# Deep Phenomics of Pattern Recognition Receptor Agonist Specific Activation of Human Blood

**DOI:** 10.1101/2025.05.12.653280

**Authors:** Soumik Barman, Arsh Patel, Aisling Kelly, David J. Dowling

## Abstract

Human whole blood (WB) immunophenotyping may represent the *in vivo* immunological state with better fidelity than artificially isolated peripheral blood mononuclear cells (PBMCs). We used a deep phenomics modeling approach to elucidate the quantitative differences in major immune cell lineages in WB and PBMC compartments in a steady-state *in vitro* setting. We studied functional innate immune responses induced by pattern recognition receptor agonist adjuvants (PRRa). Using an optimized 49-parameter CyTOF panel and implementing machine learning (ML) algorithms, we mapped PRRa-mediated CD69, CD40, CD80, CD86 and CCR7 activation at the nodal innate immune subsets level. We also portrayed cellular origin of innate functional chemokine CCL4 and intracellular cytokine IFNγ production. We mapped neutrophils as the primary source of TLR7/8 agonist (TLR7/8a) and STING agonist mediated CCL4 responses in WB. Notably, in the PBMC fraction, where neutrophils are limited, natural killer (NK) cells became the major source of innate CCL4 production. TLR7/8a-mediated IFNγ induction by early NK cells was mapped in PBMCs, which was limited in WB. Considering such distinctions, we hypothesized that deep phenomics employing a clinical sample that has not been manipulated, i.e., WB, may be additive in translating *in vitro* innate fingerprinting into *in vivo* biology.

## 1. Introduction

Advancements in phenomics, such as mass and spectral flow cytometry, along with extended usage of machine learning (ML) algorithms, have re-energized the phenotypic study of immune cells during the onset of disease or during the testing of new immune therapies and vaccines. During the COVID-19 pandemic mass cytometry (CyTOF) was used extensively to study the immune response ^[1–4]^ and vaccine efficacy ^[5–6]^ in humans. In response to the need for innovative solutions to profound scientific inquiries, several CyTOF panels were validated and established to provide a deeper understanding of the human immune system ^[7–8]^, facilitate the study of non-human primate’s leukocytes ^[9–10]^ and expand the applicability of existing murine models ^[11–12]^. Cloud-based analytical tools (e.g., the Cytobank and OMIQ platforms) have further revolutionized high-dimensional phenotypical data analysis ^[13–14]^. Automated gating strategies ^[15]^ and unsupervised clustering analysis facilitated the exploration of novel cellular phenotypes in greater depth ^[16]^. Despite these advancements, there remains little discussion and exploration on how investigators should model the *in vivo* immune response *in vitro* ^[17–18]^.

Human peripheral blood mononuclear cells (PBMCs) are widely used biological specimen to model *in vivo* biology *in vitro*. During PBMC isolation using density gradient centrifugation from whole blood (WB), granulocytes settle on top of red blood cells, while the PBMC fraction localizes between the plasma and Ficoll-Paque interface ^[19–20]^. Consequently, a significant portion of high-density neutrophils ^[21]^ is lost from PBMCs. Therefore, PBMCs do not fully mimic the *in vivo* condition ^[18, 22]^ especially due to the absence of granulocytes and enrichment of T cells along with B cells, NK cells, γδ T cells and MAIT/NKT cells. Interestingly, such enrichment makes PBMCs a better biological sample for immunophenotyping of antigen-specific rare T cells by tetramer staining ^[23]^. Phenotypic discrepancies between fresh blood and PBMCs ^[24]^ or cryopreserved PBMCs after intracellular staining ^[23]^ have been well documented. Stimulated WB exhibited a reduction in nonspecific biological noise in comparison to stimulated PBMCs ^[25]^. In general, the use of WB or PBMCs as biological samples for immunoprofiling assays may necessitate the standardization of markers of interest to mitigate such potential pitfalls ^[26]^.

Adjuvant-centric vaccine development is a growing field, with distinct focuses across basic and translational immunological research ^[27]^. Our team has helped pioneer both the phenotypic-based discovery of adjuvants ^[28]^ and the use of deep immunophenotyping of adjuvant-induced innate responses across the human lifespan ^[29]^. We have recently applied CyTOF in combination with computational tools ^[29]^ to reveal distinct, multilayered immunity in neonates, adults and elders and found that i) IFNγ responses correlate with pattern recognition receptor agonist adjuvants (PRRa), such as TLR7/8 adjuvanticity *in vitro*, ii) R848, a TLR7/8 agonist (TLR7/8a) induced robust IFNγ responses in PBMCs, which was derived predominantly from NK cells; iii) γδ T cells in adult humans contributed significantly to early IFNγ responses compared to newborn and elder; iv) other adjuvants (alum, MPLA, and CpG) were found to be inferior in eliciting IFNγ responses in all age groups. Therefore, a) innate immune responses and the adaptive immunity they facilitate, are adjuvant specific, and b) as we have demonstrated by elucidating IFNγ induction as an exemplary biomarker for effective vaccine adjuvant, deep immunophenotyping may aid in identifying the adjuvant-specific responses that can guide the development of next-generation vaccine formulations.

We now hypothesize that deep immunophenotyping of human WB, via an updated and standardized 49-parameter CyTOF panel along with adjuvant-induced innate immune mapping, will advance our understanding of driving effective adaptive responses to vaccination in human populations, by providing fresh insight into which phenotypes should be the basis for future study. To get an idea of how the baseline differences of immune cell lineages in WB and PBMCs shape the adjuvant specific innate immune profile, we used model adjuvants TLR7/8a R848 and STING agonist (STINGa) 2′3′-cGAMP, stimulated WB and PBMCs individually and captured the innate fingerprinting by mass cytometry and ML algorithms. Furthermore, our current study aimed to elucidate the PRRa-induced innate immune profile between WB and PBMC *in vitro*, thereby establishing a comprehensive *in vivo* snapshot of the human immune response.

## 2. Results

### 2.1. Panel optimization and CyTOF-based profiling among canonical immune cell lineages between WB and PBMCs

WB offers proximity towards the *in vivo* platform ^[18, 30]^, whereas PBMCs lack the abundance of granulocyte populations ^[18, 30–31]^. To elucidate the compositional disparities between innate and adaptive immune cell populations in freshly isolated WB and PBMCs, we conducted immunophenotyping of both compartments employing a 49-parameter CyTOF panel (Table 1). We used the pre-optimized Maxpar^®^ Direct^™^ Immune Profile Assay or MDIPA (Table S1) as a backbone panel, comprising a lyophilized mixture of metal-labeled 30 surface markers and rhodium as a viability marker ^[2, 32–33]^. Several clinical studies ^[34–36]^ have utilized the MDIPA due to its comprehensive nature and convenient immunophenotyping screening capabilities in a single assay. To minimize intersample variability ^[37]^ resulting from varying PRRa stimulations and acquisition time ^[38]^, we implemented a four-choose-three barcoding format (Figure S1A) utilizing anti-CD45 Live Cell Barcoding (LCB) strategy which is compatible with MDIPA ^[39]^. Re-validation of LCBs was conducted through titrations (Figure S2) utilizing freshly isolated WB and PBMCs. We have expanded the MDIPA backbone by combining it with the Maxpar^®^ Direct^™^ T Cell Activation Expansion Panel (Table S1) through specific modifications as described in the “Experimental Section”. We further validated custom conjugated Abs by titration (Figure S2) using stimulated WB and PBMCs and incorporated them into our optimized 49-parameter panel (Table 1).

**Table 1.** The standardized 49-parameter CyTOF panel for immunophenotyping of human WB and PBMCs. The Maxpar^®^ Direct^™^ Immune Profiling Assay Kit (Vendor: Standard BioTools; #201334) or MDIPA backbone (#S10310) was used for mass cytometry assay. Drop-in barcodes were highlighted in yellow color whereas drop-in surface Abs were in green and drop-in intracellular Abs were highlighted in red color.

| Index | Specificity | Clone | Metal | Purpose | Identifier | Analysis<sup>#</sup> |
| --- | --- | --- | --- | --- | --- | --- |
| 1 | CD45 | HI30 | 89Y | Leukocytes | S10310 | G, HD |
| 2 | Viability | --- | 103Rh | Exclusion | S10310 | G |
| 3 | CD40 | 5C3 | 106Cd | Activation marker | Cust* | HD, C |
| 4 | CD45 | HI30 | 110Cd | Barcode | 3110001B | G |
| 5 | CD45 | HI30 | 111Cd | Barcode | 3111001B | G |
| 6 | IL-2 | MQ1-17H12 | 112Cd | Functional cytokine | 3112002B | G, HD |
| 7 | CD69 | FN50 | 113Cd | Activation marker | 3113002B | HD, C |
| 8 | TNFα | Mab11 | 114Cd | Functional cytokine | 3114002B | G, HD |
| 9 | IFNγ | B27 | 116Cd | Functional cytokine | 3116002B | G, HD |
| 10 | CD196/CCR6 | G034E3 | 141Pr | Th17 marker | S10310 | HD |
| 11 | IL-4 | MP4-25D2 | 142Nd | Functional cytokine | 3142002B | G, HD |
| 12 | CD123 | 6H6 | 143Nd | pDCs, Basophils | S10310 | G, HD, C |
| 13 | CD19 | HIB19 | 144Nd | B cells | S10310 | G, HD |
| 14 | CD4 | RPA-T4 | 145Nd | CD4 T cell lineage | S10310 | G, HD |
| 15 | CD8a | RPA-T8 | 146Nd | CD8 T cell lineage | S10310 | G, HD |
| 16 | CD11c | Bu15 | 147Sm | mDCs | S10310 | G, HD, C |
| 17 | CD16 | 3G8 | 148Nd | Monocyte / NK differentiation | S10310 | G, HD |
| 18 | CD45RO | UCHL1 | 149Sm | Activation marker | S10310 | G, HD |
| 19 | CD45RA | HI100 | 150Nd | T cell and DC differentiation | S10310 | G, HD |
| 20 | CD161 | HP-3G10 | 151Eu | MAIT cells, ILCs | S10310 | G, HD |
| 21 | CD194/CCR4 | L291H4 | 152Sm | Th2 marker | S10310 | G, HD |
| 22 | CD25 | BC96 | 153Eu | Regulatory T cells (Treg) | S10310 | G, HD |
| 23 | CD27 | O323 | 154Sm | T and B cell differentiation | S10310 | G, HD |
| 24 | CD57 | HNK-1 | 155Gd | Senescence | S10310 | G, HD |
| 25 | CD183/CXCR3 | G025H7 | 156Gd | Th1 marker | S10310 | HD |
| 26 | CD185/CXCR5 | J252D4 | 158Gd | T<sub>FH</sub> marker | S10310 | HD |
| 27 | CD28 | CD28.2 | 160Gd | T cell memory | S10310 | G, HD |
| 28 | CD38 | HB-7 | 161Dy | Activation marker | S10310 | G, HD |
| 29 | CD80 | 2D10.4 | 162Dy | Activation marker | 3162010B | HD, C |
| 30 | CD56 | NCAM16.2 | 163Dy | Pan NK/ $\gamma\delta$ T cell activation | S10310 | G, HD |
| 31 | TCRgd | B1 | 164Dy | Pan $\gamma\delta$ T cell | S10310 | G, HD |
| 32 | CCL4 | D21-1351 | 165Ho | Functional cytokine | Cust* | G, HD |
| 33 | CD294 | BM16 | 166Er | Eosinophils | S10310 | G, HD |
| 34 | CD197/CCR7 | G043H7 | 167Er | T cell memory | S10310 | G, HD, C |
| 35 | CD14 | 63D3 | 168Er | Monocytes | S10310 | G, HD |
| 36 | IL-17A | BL-168 | 169Tm | Functional cytokine | 3169006B | G, HD |
| 37 | CD3 | UCHT1 | 170Er | T cell lineage | S10310 | G, HD |
| 38 | CD20 | 2H7 | 171Yb | B cells | S10310 | G, HD |
| 39 | CD66b | G10F5 | 172Yb | Neutrophils | S10310 | G, HD |
| 40 | HLA-DR | LN3 | 173Yb | Activation marker | S10310 | G, HD, C |
| 41 | IgD | IA6-2 | 174Yb | B cell differentiation | S10310 | HD |
| 42 | CD127 | A019D5 | 176Yb | Treg | S10310 | G, HD |
| 43 | Cell-ID Intercalator | --- | 191Ir | DNA content | S00093 | G |
| 44 | Cell-ID Intercalator | --- | 193Ir | DNA content | S00093 | G |
| 45 | CD45 | HI30 | 194Pt | Barcode | 3194001B | G |
| 46 | CD45 | HI30 | 195Pt | Barcode | 3195001B | G |
| 47 | Perforin | B-D48 | 196Pt | Functional cytokine | 3196002B | G, HD |
| 48 | Granzyme B | Gb11 | 198Pt | Functional cytokine | 3198002B | G, HD |
| 49 | CD86 | IT2.2 | 209Bi | Activation marker | Cust* | HD, C |

\*Custom conjugated antibodies (Vendor: Standard BioTools) were indicated by Cust.
<sup>#</sup>Computational analysis was done by the cloud-based platform OMIQ using cell-subset gating (G) manually followed by high-dimensionality (HD) reduction analysis using FIt-SNE and supervised clustering analysis by CITRUS (C).

After the CyTOF run, quality controlled and assured (Figure S1 & Table S2) data were analyzed by the cloud-based platform OMIQ ^[40]^. Using manual gating (Figures 1A & S3), we defined major innate and adaptive immune cell populations in both WB and PBMC compartments and portrayed the cellular topography in bivariate plots (Figures 1B) using Fast interpolation-based t-distributed Stochastic Neighbor Embedding (FIt-SNE) ^[41]^. We evaluated the behavior of FIt-SNE along with two other frequently used Dimensionality Reduction (DR) algorithms: Optimized t-SNE (opt-SNE) ^[40]^ and t-SNE that uses Compute Unified Device Architecture (CUDA)-enabled Graphics Processing Units (GPUs) to speed up the computation (t-SNE-CUDA). Although t-SNE-CUDA is the fastest algorithm ^[42]^, it was unable to preserve the coordinates of the cellular islands in the bivariate plot between subsequent runs (Figure S4A). This is likely due to the algorithm’s inability to accept a seed input. In contrast, both opt-SNE and FIt-SNE could preserve the cellular topography between subsequent runs (Figure S4B & C). We decided to use FIt-SNE algorithms for the visualization because it is 1.65 times faster than opt-SNE (Figure S4B & C).

**Figure 1.**
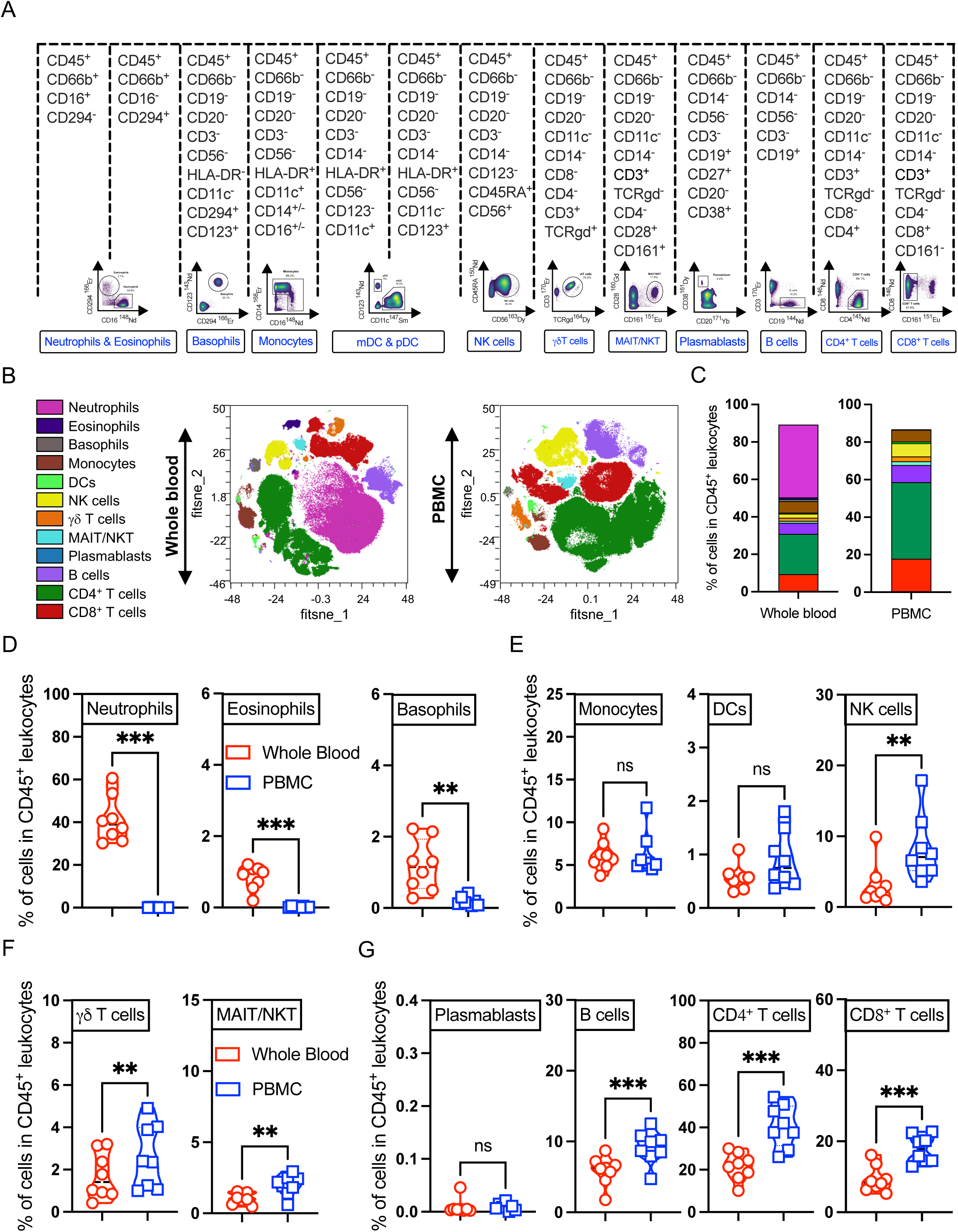
Gating strategy implementation using the 49-parameter CyTOF panel on freshly isolated human WB and PBMCs (A) portraying canonical cell lineages. Manual gating was accomplished by the OMIQ platform. B) 94K events of live CD45^+^ cells from each donor used in concatenated visualization with FIt-SNE. The FIt-SNE overlays comparing the topography of cellular landscape of WB or in PBMCs in steady-state. C) Comparison of major immune cell lineage’s frequency (as a percentage of live CD45^+^ leukocytes) in unstimulated WB or PBMCs. Such profiling reveals substantial baseline disparities between granulocyte (CD45^+^ CD66b^+^) and T cell compartments in unstimulated WB and PBMCs. D) Truncated violin plots showing the abundance of granulocytes in WB compared to PBMCs. E) Limited baseline differences in monocyte and DC compartment whereas innate NK cells, (F) γδ T cells and MAIT/NKT cells were found to be accumulated in PBMCs along with (G) B cells and T cells, when compared with WB. Statistical comparison was performed using either two-tailed paired t-test or Wilcoxon test corrected for multiple comparisons; **p < 0.01, ***p < 0.001, ns denoted non-significant. Each circle or square represents a single participant (n = 8 per group).

A significant limitation of human PBMCs is the absence of granulocyte populations ^[18, 20, 31]^. Using the OMIQ platform and DR analysis using FIt-SNE, we profiled the abundance of granulocytes (neutrophils, eosinophils and basophils) in WB compartment compared to PBMCs from a cohort of 8 healthy human adult donors (Figures 1B-D, S5A-B & Table S3). PBMCs were enriched with NK (Figures 1E & Table S3), γδ T and MAIT/NKT (Figure 1F & Table S3) cells which are the component of the innate arm. Monocytes, DCs (Figure 1E & Table S3) and plasmablasts (Figure 1G & Table S3) showed limited baseline differences. PBMCs were also found to be enriched with B cells, CD4^+^ T cells, CD8^+^ T cells (Figure 1G & Table S3) and Regulatory T (Treg) cells (Table S3) when compared with WB. WB is advantageous for studying delicate cells like neutrophils and DCs because its freshness preserves their functionality and structure, which can be lost when cells are separated and stored ^[31, 43]^.

### 2.2. PRRa-mediated innate immune response in WB and PBMCs

Innate immunoprofiling following PRRa-stimulation using fresh WB may exhibit enhanced activation and in some context, may provide a more accurate representation of the *in vivo* immunological state compared to PBMCs ^[18]^. Very recently, we portrayed R848-specific CD40 upregulation on DCs and monocytes across the human lifespan using a first-generation 37-parameter CyTOF panel using freshly isolated PBMCs ^[29]^. Here, we took advantage of both adult WB and PBMCs to investigate the phenotypic alterations by capturing the innate CD69, CD40, CD80, CD86 and CCR7 activation profile mediated by PRRa stimulation using the second-generation 49-parameter CyTOF panel.

We observed R848-mediated upregulation of CD69 on NK cells, DCs, γδ T cells, and MAIT/NKT cells from both WB and PBMCs (Figure 2), an observation that is consistent with a previous report ^[44]^. 2′3′-cGAMP exhibited a similar trend, as well as it induced CD69 upregulation in monocytes from PBMC only. Such upregulation activates lymphocytes or NK cells ^[44]^. R848 showed superiority in CD40 activation on DCs in both WB and PBMCs compared with 2′3′-cGAMP (Figure 3A & B). R848 also upregulated CD80 expression on monocytes and DCs in both WB and PBMCs (Figures 3C & D). Interestingly, 2′3′-cGAMP induced CD40 and CD80 on monocytes in both WB and PBMCs (Figure 3A-D) while CD80 was upregulated on DCs in WB only (Figure 3C). WB showed upregulation of CD86 on DCs without stimulation (Figure 3E) despite in PBMCs compartment we found R848 meditated CD86 upregulation on DCs (Figure 3F) whereas 2′3′-cGAMP induced CD86 only on monocytes. Such upregulation of CD40/CD80/CD86 initiate the activation of DCs which ultimately initiate T cell priming ^[29]^ as well as T cell proliferation ^[44]^. We did not observe any upregulation of CCR7 in the monocyte compartment of PBMC (Figure 3H), despite we did see upregulation in the monocyte compartment of WB in steady-state (Figure 3G). R848-mediated CCR7 upregulation was captured in the DC compartment in both WB and PBMCs (Figures 3G & H). Therefore, TLR7/8a R848 has the potential to coordinate the migration of DCs via CCR7 to secondary lymphoid organs like lymph nodes, to initiate adaptive immune responses ^[45–46]^.

**Figure 2.**
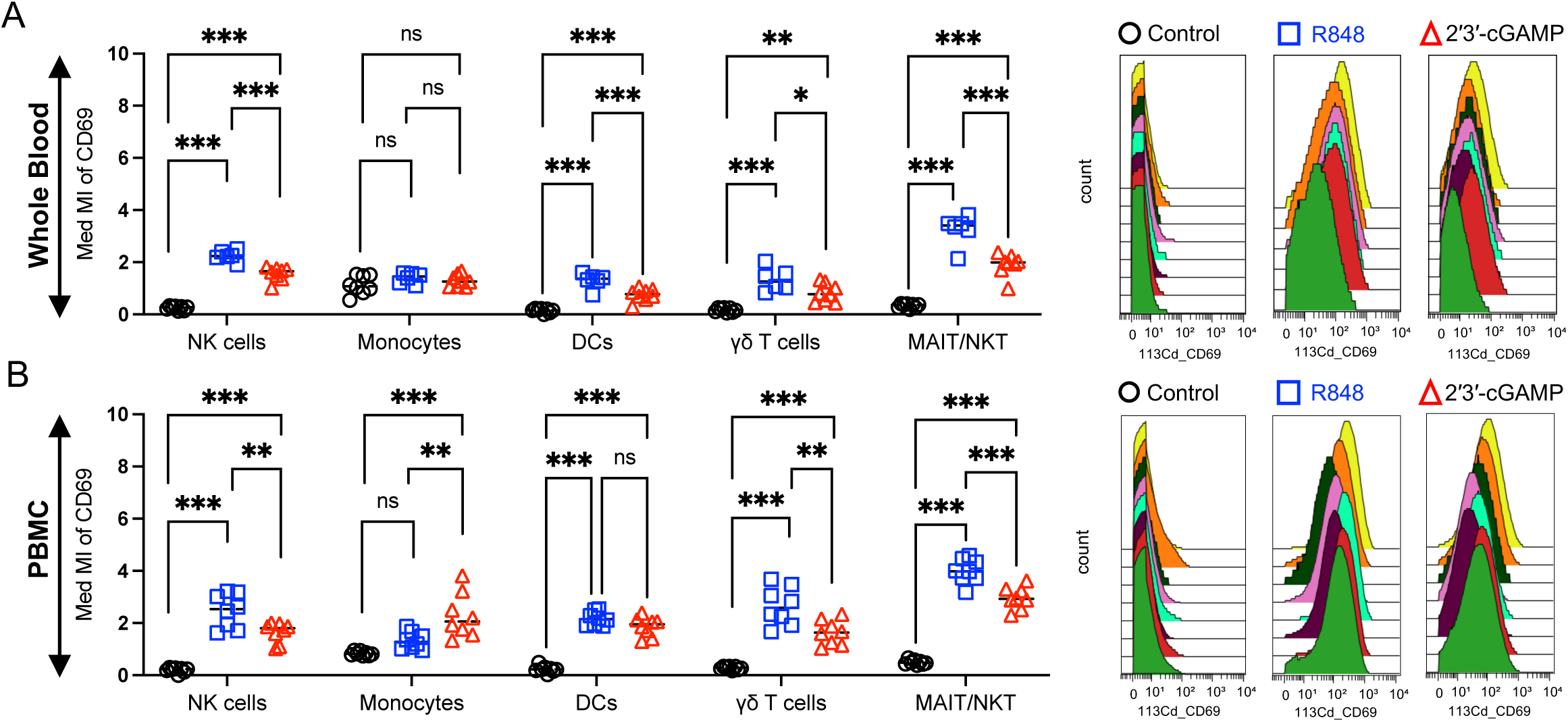
TLR7/8a R848 (5 μM) and STINGa 2′3′-cGAMP (25 μg/ml) induce upregulation of CD69 on NK cells, DCs, *γδ* T cells and MAIT/NKT cells in both (A) WB and in (B) PBMCs. Representative median metal intensity (Med MI) of CD69 is also shown in histogram (to the right of each row) on MAIT/NKT cells in both WB and in PBMCs. Color in the histogram indicates each participant’s (n = 6-8) CD69 activation profile upon PRRa stimulation. Two FCS files (R848 stimulated WB) are excluded from the analysis as they do not possess 94K events of live CD45^+^ population during “Subsampling” as described in the “Experimental Section”. Statistical comparison was performed using either one-way ANOVA or nonparametric Kruskal-Wallis test corrected for multiple comparisons; *p < 0.05, **p < 0.01, ***p < 0.001, ns denoted non-significant. Each circle, square or triangle represents a single participant (n = 6 – 8 per group).

**Figure 3.**
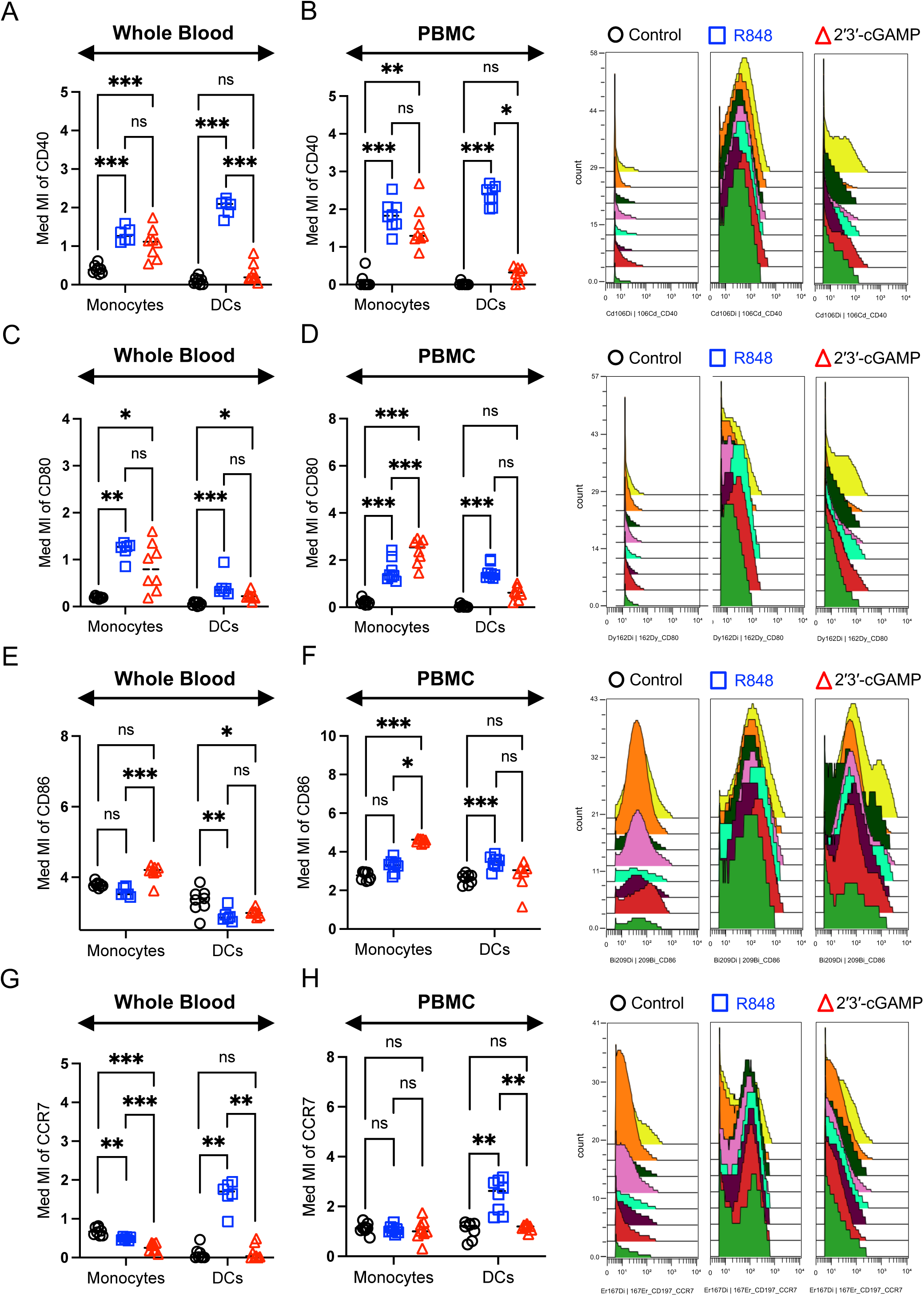
TLR7/8a R848 drives CD40, CD80, CD86 and CCR7 upregulation on DCs in PBMCs. Activation markers (A, B) CD40; (C, D) CD80; (E, F) CD86 and (G, H) chemokine receptor CCR7 were evaluated in both monocytes and DCs either in (A, C, E, G) WB or in (B, D, F, H) PBMC respectively after stimulations. Representative median metal intensity (Med MI) of CD40, CD80, CD86 and CCR7 expression on DCs in PBMCs is shown to the right of each row. Color in the histogram indicates each participant’s CD40/CD80/CD86/CCR7 activation profile upon PRRa stimulation either with R848 (5 μM) or 2′3′-cGAMP (25 μg/ml). Statistical comparison was performed using either one-way ANOVA or nonparametric Kruskal-Wallis test corrected for multiple comparisons; *p < 0.05, **p < 0.01, ***p < 0.001, ns denoted non-significant. Each circle, square or triangle represents a single participant (n = 6 – 8 per group). Two FCS files (R848 stimulated WB) are excluded from the analysis because they lack 94K events of the live CD45^+^ population during the “Subsampling” process, as described in the “Experimental Section”.

### 2.3. Dissecting the stratifying signatures of pDC clusters in R848 stimulated PBMCs by CITRUS algorithm

After portraying R848 and 2′3′-cGAMP-mediated activation profile on DCs, we tried to further dissect the DC compartment. We observed a substantial decrease in the number of myeloid dendritic cells (mDCs) in both adult WB and PBMCs following R848 stimulation compared to both control and 2′3′-cGAMP-treated groups (Figures 4A & B). R848 exclusively increased the frequency of pDCs in PBMCs (Figure 4B), which was absent in WB (Figure 4A) following R848 stimulation.

**Figure 4.**
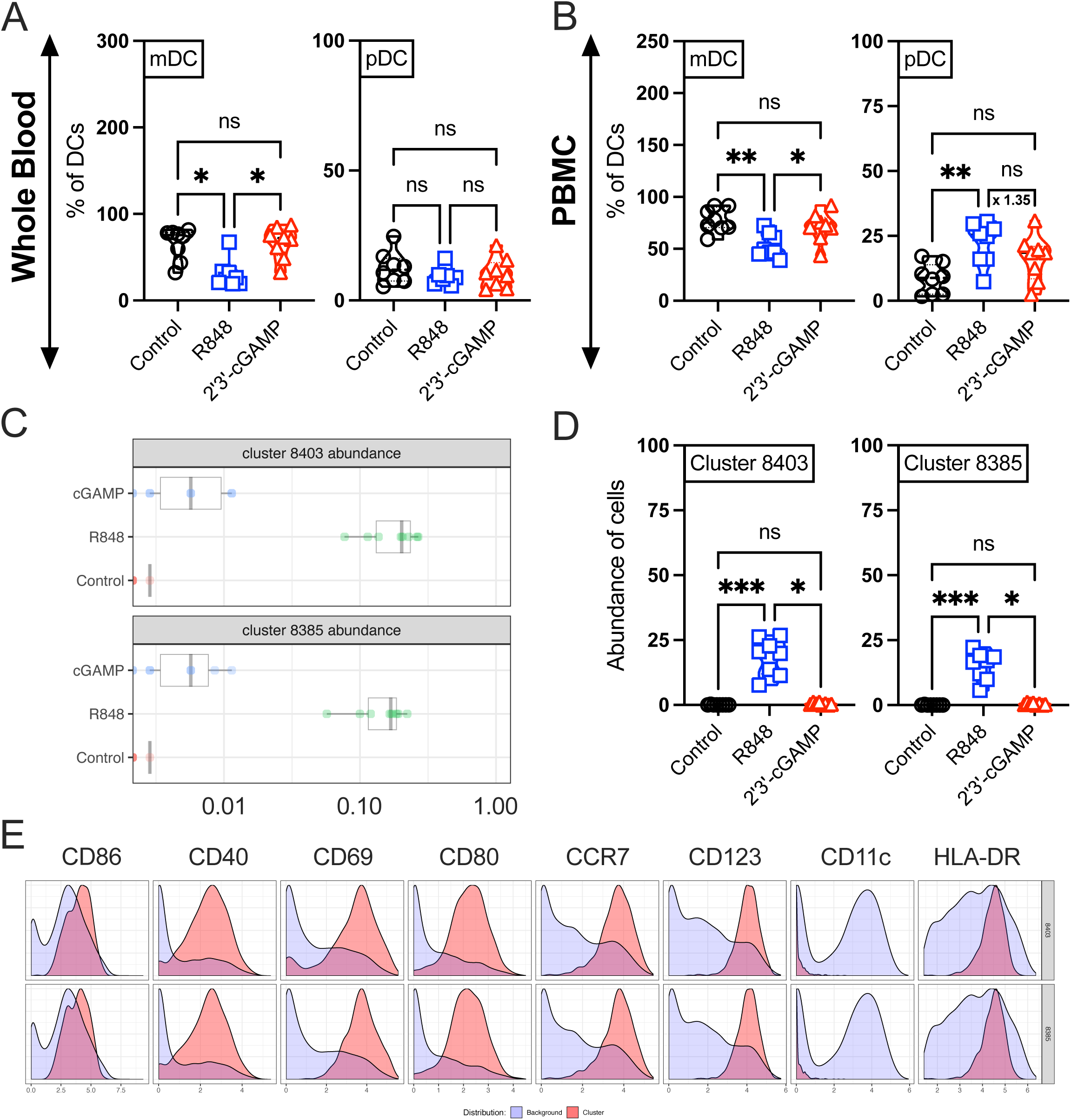
TLR7/8a R848 drives accumulation and activation of pDCs in PBMCs. The frequency of DC subpopulations in (A) WB and (B) PBMC after R848 (5 μM) and 2′3′-cGAMP (25 μg/ml) stimulation after “Subsampling” as described in the “Experimental Section”. R848 maintained pDC accumulation (B) in PBMC after stimulation. Mean fold difference between R848 and 2′3′-C) Supervised clustering analysis by CITRUS using manually gated DCs in PBMCs predicted quantitative abundance of most stratifying pDC clusters (8403 & 8385) between groups. D) Significant abundance of targeted clusters from (C) were validated manually by the OMIQ platform using the categorical filters for downstream analysis and presented as a percentage of DCs. Phenotypic expression of these clusters is also shown in (E). The expression level of each selected marker was represented by a cluster (red) over its background (light blue). Figures were directly exported from the OMIQ platform for C & E. Statistical comparison was performed using either one-way ANOVA or nonparametric Kruskal-Wallis test corrected for multiple comparisons; *p < 0.05, **p < 0.01, ***p < 0.001, ns denoted non-significant. Each circle, square or triangle represents a single participant (n = 8 per group). Prediction Analysis for Microarrays (PAMR) regression model with a threshold of 1% False Discovery Rate (FDR) employed by CITRUS as a statistical methodology to identify the stratifying signatures among distinct subpopulations of DC as described in “Experimental Section”.

Next, we employed the ML algorithm CITRUS (cluster identification, characterization, and regression) via the OMIQ platform to further characterize functional plasticity ^[47]^ induced by PRRa in the pDC compartment. CITRUS identifies cell populations associated with a specific biological or clinical parameter ^[48]^. Additionally, CITRUS incorporates regularized regression modeling to ensure statistically robust comparisons between groups ^[13]^, which is particularly important due to the heterogeneity of any clinical cohorts ^[49]^. To identify such stratifying signatures in the pDC compartment, we initially studied PRRa-stimulated PBMCs using CITRUS, which identified 32 clusters (Figure S6A) distributed in three distinct islands. Based on the expression of CD11c (Figure S6C) and CD123 (Figure S6D), we designated the islands as mDC island, mDC predominant island, and pDC island (Figure S6A). Notably, in the mDC predominant island, all clusters exhibited mDC-associated phenotypes, except for clusters 8403 and 8385 (Figures S6A, C & D). CITRUS identified a substantial abundance of clusters 8403 and 8385 in the R848 stimulated pDC compartment (Figures 4C & D). The parent pDC cluster 8403 and its offspring cluster 8385 exhibited highly activated identical phenotypes, characterized by the upregulation of CCR7, CD80, CD69, CD40, and CD86 (Figure 4E).

We also mapped the stratifying signatures of clusters 8416 and 8405, which belong to the pDC island (Figure S6A). The pDC clusters 8416 and 8405, which were abundant in 2′3′-cGAMP-stimulated PBMCs (Figures S6E and F), exhibited upregulation of both CD80 and CD69 (Figure S6G). CD40 was upregulated in the offspring cluster 8405, while it was lower in the parent cluster 8416 (Figure S6G). Stimulated pDCs by STINGa likely remain non-migratory in nature due to their low CCR7 induction (Figure S6G). STINGa adjuvantation and its antitumor therapeutic efficacy via DCs has been explored in murine models ^[50–51]^. TLR7a-specific DC accumulation and migration to secondary lymphoid organs were also explored in murine models ^[52–53]^. Comprehensive research examining the PRRa-mediated accumulation/activation of pDCs in PBMCs holds the potential to elucidate the mechanisms underlying the shaping of adaptive or anti-tumor immunity by these agonists.

### 2.4. The predominant cellular source of PRRa-mediated CCL4 induction is neutrophils in WB but NK cells in PBMCs

After profiling PRRa-mediated monocytes and DC activation, we focused on early cytokine/chemokine production in WB and PBMCs after 18h of PRRa-stimulation using CyTOF. R848 and 2′3′-cGAMP-mediated upregulation of intracellular cytokines (IL-2, TNF, IL-4, IL-17) and secretory proteins (Perforin, Granzyme B) was not observed in both WB and PBMC compartments (Figure S5C & D). Interestingly, we captured R848 and 2′3′-cGAMP-mediated induction of chemokine CCL4 in WB (Figures 5A & B) and PBMCs (Figures 5E & F), respectively. In murine, MF59, CpG and alum-mediated transcriptomic signature of CCL4 was reported ^[54]^. Alum and MF59 also induce CCL4 in human immune cells ^[55]^, but the cellular sources of CCL4 have not been previously reported.

**Figure 5.**
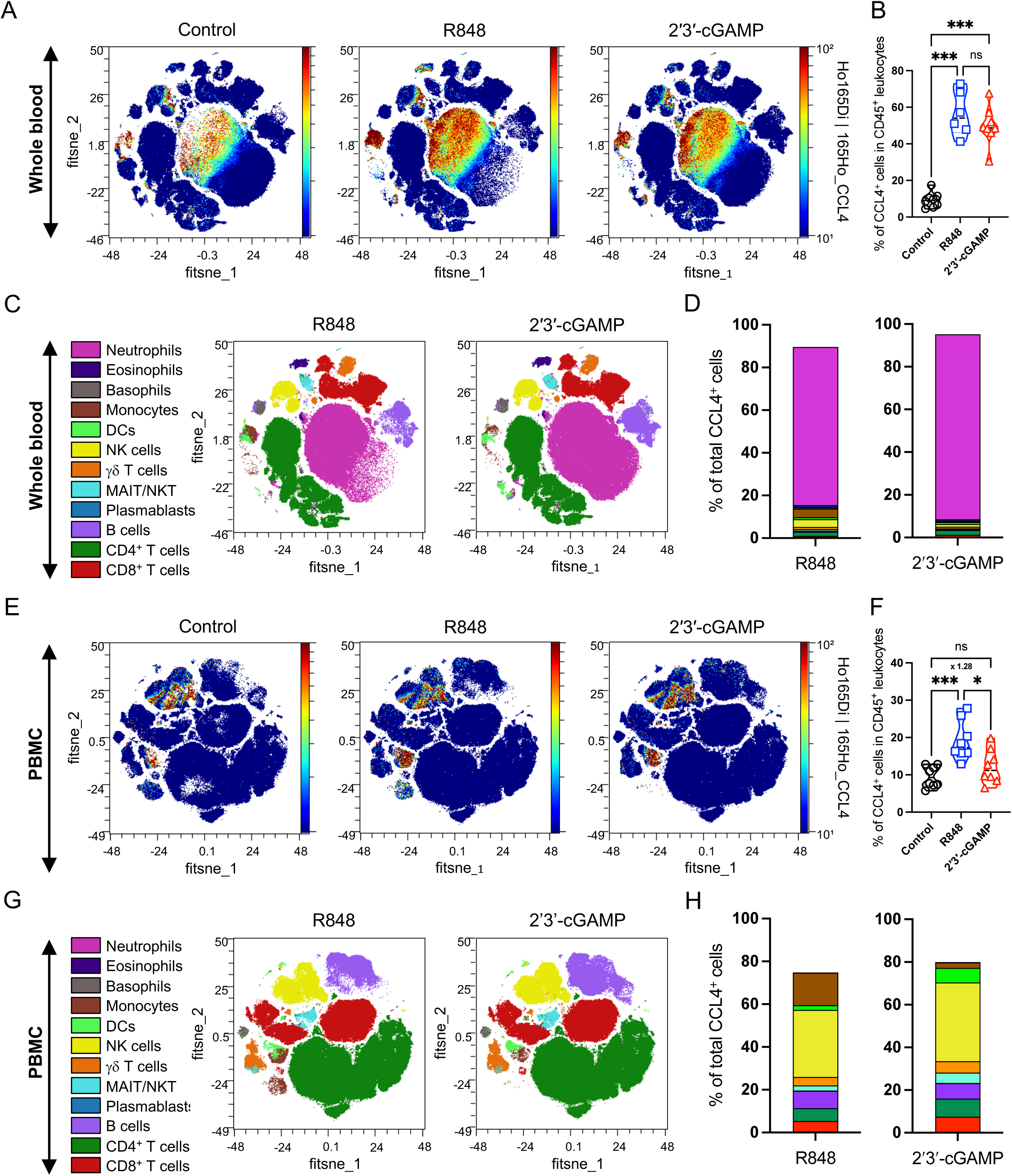
Mapping the cellular abundance of CCL4-producing leukocytes by FIt-SNE reveals a unique role of neutrophils in (A-D) WB and (E-H) NK cells Concatenated representation of CCL4 expression (in Z-axis channel) of leukocytes after R848 (5 μM) and 2′3′-cGAMP (25 μg/ml) stimulation in (A) WB and in (E) PBMCs was overlaid on FIt-SNE embedding. Color indicates CCL4 expression ranging from low (blue) to high (red). Comparison between the frequency (as a percentage of the CCL4^+^ leukocytes) in (B) WB or in (F) PBMCs after stimulation. The mean fold difference of 2′3′-cGAMP-mediated CCL4 levels relative to the steady state is also shown. High-dimensional FIt-SNE analysis of live CD45^+^ cells with overlay of 12 selected major immune cell subsets in concatenated visualization indicating (C) neutrophils in WB and (G) NK cells are the predominant source of CCL4 secretion. D, H) Comparison between the frequency (as a percentage of the CCL4^+^ leukocytes) of the major immune cell lineages after R848 and 2′3′-cGAMP stimulation to find the key source of innate CCL4 production (n = 6 - 8 per group). Statistical comparison was performed using either one-way ANOVA or nonparametric Kruskal-Wallis test corrected for multiple comparisons; *p < 0.05, ***p < 0.001, ns denoted non-significant. Each circle, square or triangle represents a single participant (n = 6 – 8 per group). Two FCS files (R848 stimulated WB) are excluded from the analysis as they do not possess 94K events of live CD45^+^ population during “Subsampling” as described in the “Experimental Section”.

After seeing the robust induction of CCL4 by R848 and 2′3′-cGAMP (Figures 5B & F), we tried to capture the predominant source of CCL4 production mediated by major innate cell lineages in WB and in PBMCs. Figure 5A & E shows the predominant island of CCL4 secreting leukocytes in FIt-SNE plots after virtual concatenation by the OMIQ platform. By FIt-SNE embedding along with virtual concatenation, we overlaid all the manually gated major innate and adaptive immune cell lineages among R848 and 2′3′-cGAMP stimulated leukocytes and visually found the CCL4-secreting predominant island belongs to the neutrophil island colored by magenta (Figure 5C) in WB. In PBMC, we found the CCL4 secreting predominant island belongs to NK cells island colored by yellow (Figure 5G). We also identified a statistically significant abundance of neutrophils (Figures 5D, S7A & B) and NK cells (Figures 5H, S7C & D) among CCL4^+^ leukocytes in PRRa-stimulated WB and in PBMCs, respectively using the OMIQ platform. We noted that CCL4 induction after 2′3′-cGAMP stimulation in PBMCs was higher than the control group (1.28-fold, Figure 5F), despite an absence of statistical significance. These data demonstrated quantitative distinctions among the primary innate chemokine-producing cellular sources based on biological samples.

We dissected the NK cell compartment (CD56^+^ subsets) according to the expression of CD57. CD56^+^ CD57^-^ cells were phenotyped as early NK cells and CD56^+^ CD57^+^ cells as late NK cells (Figure S3B). Significant abundance of CCL4 secreting early NK cells were observed in R848 stimulated PBMCs compared to late NK cells (Figure S7E). In 2′3′-cGAMP stimulated PBMCs, CCL4 secreting early NK cells were abundant (1.55-fold, Figure S7G) despite the lack of statistical significance, when compared with late NK cells. It has been documented that TLRa-mediated CCL4 secretion from freshly isolated monocytes (from PBMCs) can be synergistically enhanced by the combination of TLR4/8a and TLR3/8a ^[56]^. Here we captured monocyte-mediated CCL4 secretion in PBMC only after R848 stimulation (Figures 5H & S7C). Further immunophenotyping confirmed the classical monocyte’s involvement in TLR7/8a-mediated CCL4 secretion (Figure S7F).

### 2.5. Early NK cell-mediated substantial IFNγ responses were captured in R848 stimulated PBMCs but not in WB

Recently, we mapped NK cells as the key source of TLR7/8a specific innate IFNγ responses in PBMCs across immunologically distinct age groups ^[29]^. Using the 49-parameter CyTOF panel here, we additionally captured R848 and 2′3′-cGAMP-mediated induction of IFNγ in both WB (Figure 6A & B) and PBMCs (Figure 6C-G). PRRa-stimulated WB did not demonstrate any baseline differences in IFNγ induction between the steady and stimulated states (Figure 6B). In PBMCs, we observed substantial IFNγ secretion induced by R848 among live cells (Figure 6D). In consistent with our previous study ^[29]^, we profiled NK cells as the predominant source of IFNγ secretion in R848-stimulated PBMCs visually by FIt-SNE embedding along with virtual concatenation (Figures 6C & E). We also observed a significant increase in the number of IFNγ^+^ NK cells in the live CD45^+^ IFNγ^+^ compartment following R848 stimulation (Figure 6F & G). Comprehensive immunophenotyping of the IFNγ^+^ NK cell compartment further confirmed the involvement of early NK cells in TLR7/8a R848-mediated IFNγ secretion (Figure S7H). We also observed R848-mediated IFNγ secretion in the γδ T cell and B cell compartments (Figure 6G), but these findings were not statistically significant compared to other targeted cell populations.

**Figure 6.**
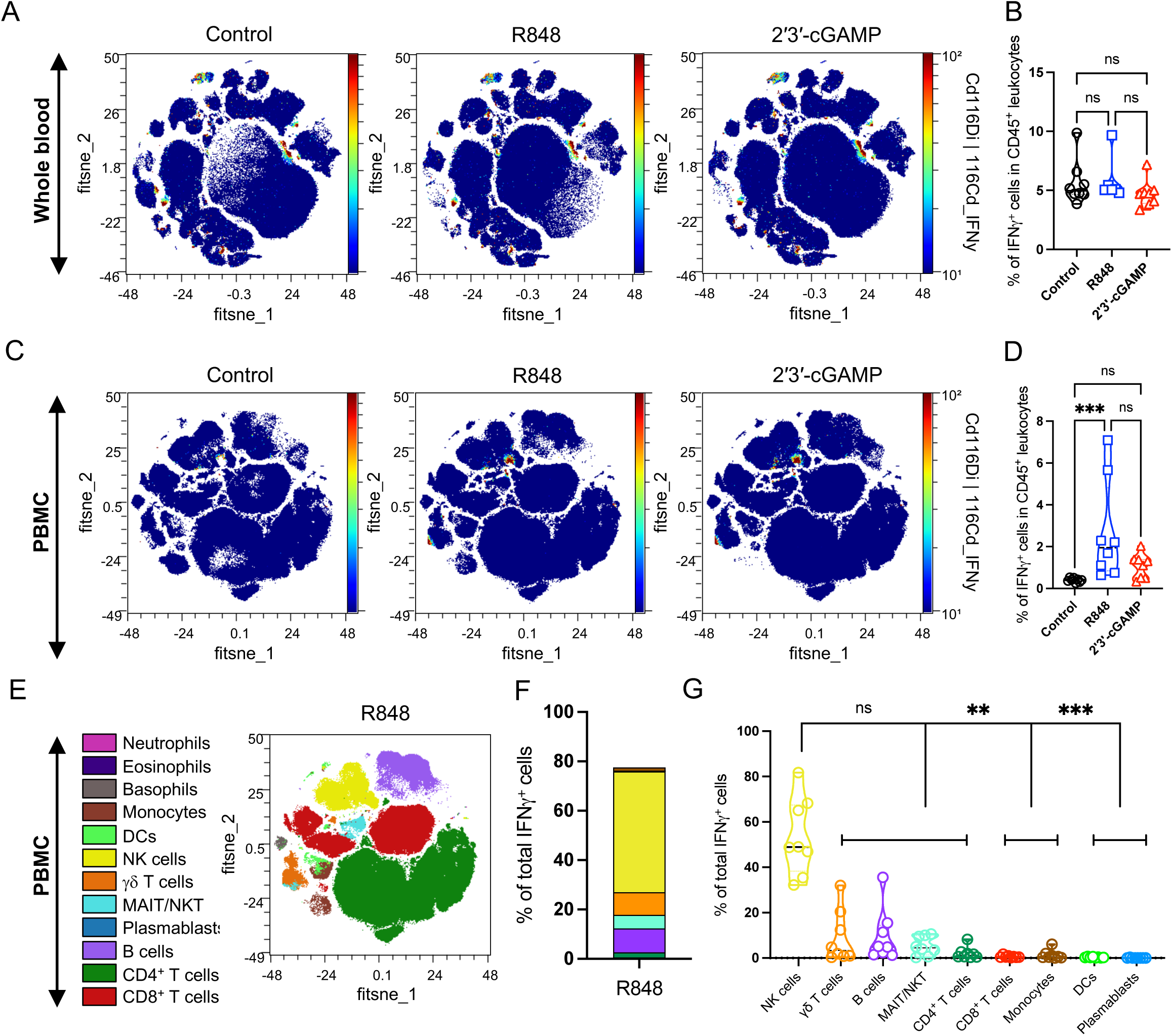
Mapping the cellular abundance of IFNγ producing leukocytes using FIt-SNE algorithm reveals a unique role of NK cells in (C-G) PBMCs. Concatenated representation of IFNγ expression (in Z-axis channel) of leukocytes after R848 (5 μM) and 2′3′-cGAMP (25 μg/ml) stimulation in (A) WB and in (C) PBMCs was overlaid on FIt-SNE embedding. The color scale indicates IFNγ expression ranging from low (blue) to high (red). Comparison between the frequency (as a percentage of the IFNγ^+^ leukocytes) in (B) WB or in (D) PBMCs after stimulation. High-dimensional FIt-SNE analysis of live CD45^+^ cells with overlay of 12 selected major immune cell subsets in concatenated visualization indicating (E) NK cells are the predominant source of IFNγ secretion. F, G) Comparison between the frequency (as a percentage of the IFNγ^+^ leukocytes) of the major immune cell lineages after R848 stimulation to find the key source of innate IFNγ production (n = 8 per group). Statistical comparison was performed using either one-way ANOVA or nonparametric Kruskal-Wallis test corrected for multiple comparisons; **p < 0.01, ***p < 0.001, ns denoted non-significant. Each circle, square or triangle represents a single participant (n = 8 per group). Two FCS files (R848 stimulated WB) are excluded from the analysis as they lack 94K events of live CD45^+^ population during “Subsampling” as described in the “Experimental Section”.

## 3. Discussion

The objective of this study was to evaluate the disparity in abundances of canonical immune cell lineages between the WB and PBMC compartments using deep phenomics by an optimized 49-parameter CyTOF panel. Furthermore, we investigated how PRRa as vaccine adjuvants influence innate fingerprinting within the WB and PBMC compartments. Inclusion of established activation makers, such as CD40, CD80, CD86, CD69 along with profiling innate cytokines (TNF, IFNγ, IL-2, IL-4, IL-17A), chemokine (CCL4) and exhaustion makers (Perforin, Granzyme B) allows delineation of functional immune phenotypes triggered by classical immune cell subsets (monocytes, NK cells, DCs, granulocytes) in the innate arm of human immune responses after PRRa stimulation. Using the cloud-based analytical tool the OMIQ platform and DR algorithm FIt-SNE, we portrayed baseline differences of targeted immune cell lineages in adult WB and PBMC compartments (Figure 7). We found neutrophils as the most abundant leukocytes in WB, which is limited in PBMCs and consistent with previous studies ^[57–58]^, whereas the PBMC compartment was highly enriched with CD4 and CD8^+^ T cells ^[24, 59]^.

**Figure 7.**
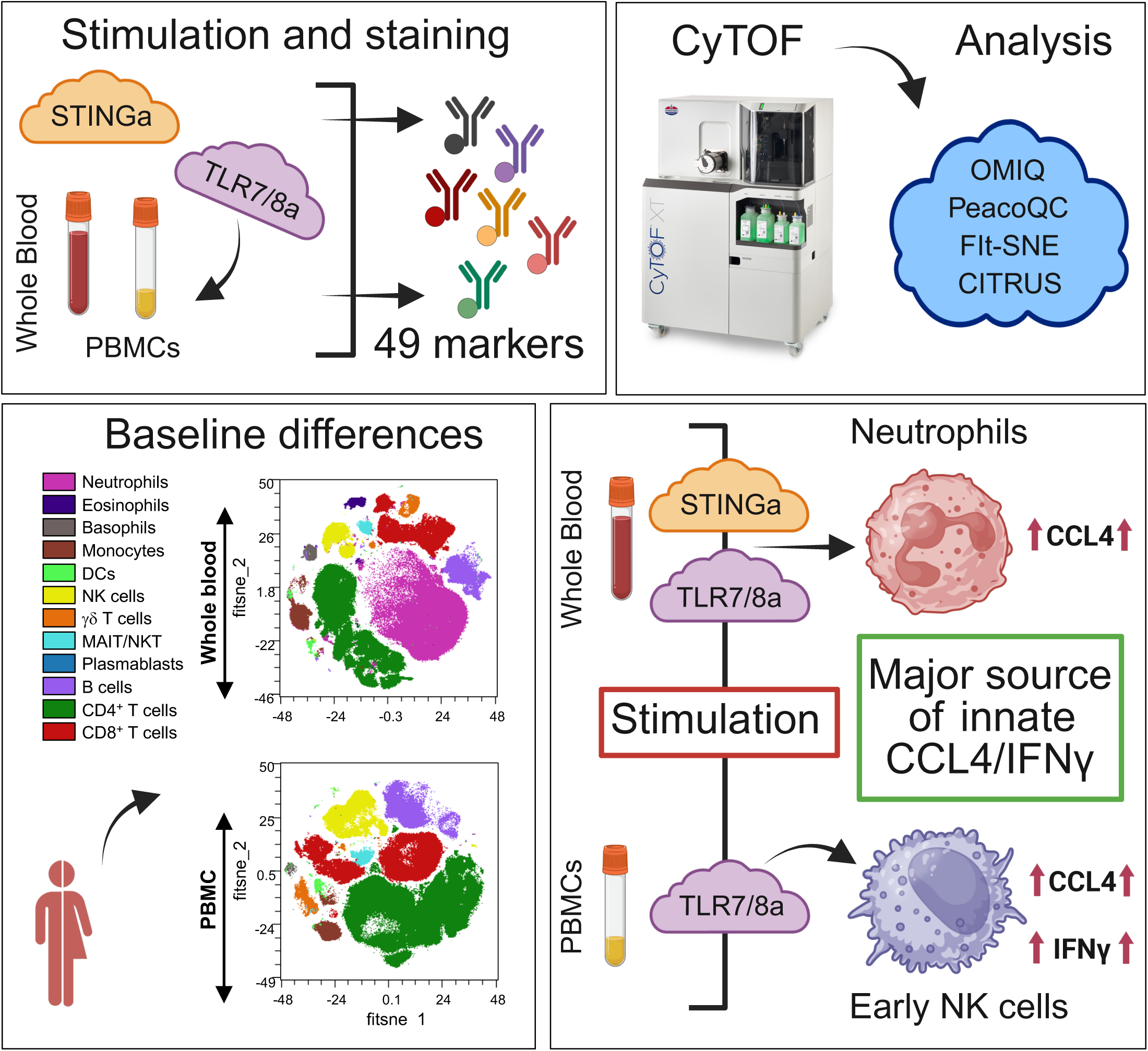
Schematic diagram illustrating the utilization of a 49-parameter CyTOF panel in conjunction with ML algorithms to elucidate the cellular origin of innate chemokine and intracellular cytokine production. Neutrophils were more abundant in WB whereas CD4^+^ T cells were more prevalent in PBMCs. Such variations in the major immune cell populations between WB and PBMCs have a notable impact on the subsequent innate chemokine/cytokine response elicited by PRRa stimulation. Therefore, for *in vitro* assays, careful consideration should be taken. [Created in BioRender. Barman, S. (2025), Agreement number: YY2893R1BQ].

Neutrophils, as the most abundant leukocytes in human blood, play critical roles in innate immunity as circulating phagocytes ^[57]^. Neutrophils also have a pivotal role in cancer progression ^[60]^ and are involved in the development of a wide range of diseases when dysfunctional (i.e. autoimmune, metabolic, genetic disorder and infectious diseases) ^[20, 61]^. Although neutrophils are the first and most abundant cell type recruited to the injection site after vaccine/adjuvant administration, they are often overlooked in traditional *in vitro* assays ^[22]^. Our data indicated that model PRRa R848 and 2′3′-cGAMP both stimulate neutrophils to robustly produce the chemoattractant CCL4 in WB (Figure 7). As a chemoattractant, CCL4 may trigger leukocyte recruitment, including NK cells ^[62]^, B cells and professional APCs ^[63]^ and initiate adaptive arm of the human immune response ^[63]^. Therefore, it is crucial to characterize how PRRa-mediated innate CCL4 response shapes adaptive immune responses in the host using clinically approved adjuvants. CCL4 can be used as an effective adjuvant in DNA vaccines designed for fish ^[64]^. CCL4 is also induced by mRNA vaccine BNT162b2 ^[65]^ or inactivated Covaxin ^[66]^ designed for SARS-CoV-2.

After portraying neutrophils as the cellular source of CCL4 secretion in PRRa-stimulated WB by using deep phenomics and ML algorithm, next we focused on PBMCs where granulocytes are limited in abundance. NK cells were identified as the CCL4-secreting cellular source after TLR7/8 and STINGa stimulation. Monocyte-mediated CCL4 secretion was also observed in PBMCs following TLR7/8a stimulation, which was found to be absent after STINGa stimulation. In the past, our group observed R848-mediated significant CCL4 accumulation in newborn WB when compared with adult WB ^[67]^. Human monocyte-specific and differentiated DC-specific CCL4 secretion was reported after alum and MF59 stimulation ^[55]^. In a follow-up study led by the same research group, using a murine model, it was found that MF59 has a stronger impact than alum on the recruitment of neutrophils at the site of administration ^[68]^. MF59, CpG and alum mediated transcriptomic signature of CCL4 in murine muscle was reported ^[54]^. Consequently, we hypothesized that for adjuvant screening, CCL4 could be an effective biomarker.

Our recent study demonstrated that early IFNγ production in adults may serve as a biomarker to predict effective adjuvanticity ^[29]^. In this study, we replicated the same trend that NK cells are the key source of R848 specific innate IFNγ responses in PBMCs. Comprehensive immunophenotyping employing the expanded 49-parameter panel corroborated the involvement of early NK cells in the TLR7/8a-specific innate IFNγ induction process (Figure 7). The absence of such phenotypes in WB raises the pertinent question of which assays are most effective in capturing *in vivo* biology. We posit that WB is the most suitable biological specimen for studying innate immune responses due to its proximity to *in vivo* biology. As PBMCs are enriched with adaptive immune cell populations, i.e. T cells, DCs, B cells, it will be wise to study adaptive immune response using deep phenomics to target rare Th1/2/17 polarization or antigen-specific T cells by tetramers.

In adjuvant biology, dendritic cells (DCs) play a crucial role by enhancing the immune response to vaccines. Adjuvants stimulate DCs, which then present antigens to T cells, leading to a stronger and more effective immune response. CD69, upregulated by PRRa, retains immune cells at the activation site, enhances cytokine production, and regulates effector functions, amplifying innate immune responses ^[69]^. We successfully captured such trends in NK cells, DCs, γδ T cells, MAIT/NKT cells in both WB and PBMCs following TLR7/8a or STINGa stimulations. Therefore, CD69 could act as an essential biomarker of adjuvant screening. We also captured PRRa-mediated activation of traditional maturation/activation markers CD40, CD80, CD86, and CCR7 and observed PRRa-specific activation differences in both DCs and monocytes, which portray how the immune cells respond to different PRRs. We re-established here the trend of TLR7/8a-mediated abundance of pDCs ^[70]^ in stimulated PBMCs. Employing the CITRUS algorithm, we comprehensively analyzed the PRRa-driven stratifying signatures of pDC subpopulations. CITRUS predicted highly activated pDC abundances in response to R848 stimulation. STINGa further contributed to the substantial activation of pDCs through the CD69/CD80 pathway. STING agonists have not yet received complete licensure as a vaccine adjuvant ^[71]^. However, they are being explored in clinical trials for cancer immunotherapy ^[72–74]^. In contrast, TLR7/8a have already been approved for clinically use due to their ability to induce Th1 responses ^[29, 75]^.

Our study features several strengths as a) our work not only portrayed the development of a 49-parameter CyTOF panel which works effectively in both WB and PBMCs in steady-state as well as in stimulated conditions, but also highlighted the application of WB or PBMC immunophenotyping assays as a predictive model ^[76]^ for adjuvant discovery and development ^[77]^; b) deep phenomics might shed light on how clinically approved adjuvants shape adaptive immunity via the CD69/CCL4/IFNγ axis as chosen predictive biomarkers for effective adjuvanticity. Furthermore, c) our standardized panel could be useful in cancer ^[78]^ and neurodegenerative disease research. Deep phenomics by CyTOF offers promising avenues for exploring uncharted territories in neurodegenerative disease research, particularly in Alzheimer’s disease (AD) ^[79]^ and dementia with Lewy bodies (DLB) ^[80]^. We hypothesize that CyTOF and ML algorithms utilizing brain autopsy ^[81]^, WB ^[82]^ or cerebrospinal fluid ^[83–84]^ can predict the cellular origin of detrimental proteins/cytokines, such as beta-amyloid ^[85]^ and tau, in AD and DLB cohorts. This prediction holds significant promise for the development of a diagnostic tool, which could subsequently pave the way for the creation of a potential immunotherapy. Finally, d) our panel can be easily modified, if researchers desire to customize it for specific needs.

Our study also has a few limitations; a) despite WB potentially providing a more accurate depiction of the *in vivo* environment, neither WB nor PBMCs may be ideal for accurately representing tissue-specific innate immune responses following adjuvanted vaccination ^[86–87]^. Implementation of our approach using 3D tissue construct ^[87]^ or lymph node fine needle aspirates ^[88]^ using clinical cohorts (from clinical trials of adjuvanted vaccines) will be more accurate in capturing the *in vivo* biology; b) we used model PRRa R848 and 2′3′-cGAMP for hypothesis generation. The same study with clinically approved adjuvants AS01B (component of license vaccines *Cervarix*, *Fendrix* and *Shingrix*), MF59 (component of license vaccine *Fluad*), and CpG-1018 (component of license vaccine *Heplisav-B*) might provide an improved rational basis for the development of next-generation adjuvantation systems tailored to enhance immunity in distinct human populations; c) there are no baseline differences in DC frequency between WB and PBMC. The concerns might arise as to how PRRa activate blood DCs? Blood DCs are phenotyped as CD141^+^ and CD1c^+^ DCs ^[89–90]^. The further incorporation of blood DC markers into our panel may facilitate the exploration of novel phenotypic characteristics in PRRa-stimulated WB.

Taken together, we hypothesized that WB could be selected as the standard *in vitro* system to facilitate the translational prediction of innate immune fingerprint, thereby mimicking the *in vivo* biology. Therefore, our immunophenotyping approach allowed us to dissect cellular functional repertoire changes which might contribute to the continued advancement of human immunology and the exploration of its variability in health and disease.

## 4. Experimental Section

### Study participants

This study involved eight individuals (4 males and 4 females) aged between 22 and 33. Blood samples were collected between August and September 2023. Sex-based analysis was not performed.

### Ethics approval and Patient consent statement

Peripheral blood from healthy, adult volunteers was collected after written informed consent with approval from the Ethics Committee of Boston Children’s Hospital, Boston, MA (protocol number X07-05-0223). All ethical regulations relevant to human research participants were followed.

### Human blood processing and *in vitro* stimulation for CyTOF

Human blood was anticoagulated with 20 units/ml pyrogen-free heparin (Table S1). All blood products were maintained at room temperature (RT) and processed or stimulated within 4h (hours) of collection. For WB stimulation, 270 µl of WB was transferred to a 48-well flat-bottom plate and diluted (1:1) with 1X T cell media (TCM) ^[91]^. For PBMCs stimulation, PBMCs were isolated from fresh WB via Ficoll gradient separation. PBMCs were also resuspended in 1X TCM and plated at a density of 6×10^6^ cells/540 μl in 48-well flat-bottom plates. Per well, 40 µl of stimulation cocktail (prepared using PRRa in 1X TCM) was added to achieve a total volume of 580 µL, which contained either WB or PBMCs. The concentration of TLR7/8a R848 (5 μM) and STINGa 2′3′-cGAMP (25 μg/ml) were determined empirically based on the results of previous studies ^[29, 92]^. After 13h of incubation in a humidified incubator at 37°C (in 5% CO_2_), Brefeldin A (1:1000) and Monensin (1:1500) (Table S1) were added to all the wells (20 μl/well) to inhibit cytokine production and facilitate optimal intracellular profiling by CyTOF. Cultures were maintained in a humidified incubator at 37°C (in 5% CO_2_) for an additional 5h.

### Mass Cytometry (CyTOF)

#### Panel design

For the development of our second-generation 49-parameter CyTOF panel, we used pre-optimized MDIPA (Standard BioTools) as a backbone panel ^[39]^. We extended the panel using Maxpar^®^ Direct^™^ T Cell Activation Expansion Panel (Standard BioTools) (Table S1), which is an optimized combination of eleven functional surface and intracellular markers ^[39]^. We replaced CD107a_106Cd, CTLA-4_162Dy, and IL-10_165Ho on the Expansion Panel with custom conjugated or drop-in Abs CD40_106Cd, CD80_162Dy, and CCL4_165Ho (Table 1). Furthermore, CD86_209 Bi was incorporated to monitor PRRa-mediated activation profiles on monocytes and DCs. For custom conjugation, Maxpar^®^ Antibody Conjugation Service with biochemical confirmation was purchased from Standard BioTools. We further validated custom conjugated Abs through titration (Figure S2) using stimulated WB and PBMCs and incorporated them into our optimized 49-parameter panel (Table 1). MDIPA-compatible four-choose-three barcoding strategy (Figure S1A) using anti-CD45 LCBs (110/111 Cd and 194/195 Pt) was adopted from ^[39]^.

#### Titrations of CyTOF Abs

For the titrations of anti-CD45 LCBs and custom-conjugated CyTOF Abs (CCL4, CD40, and CD86), WB and PBMCs were either stimulated with Mitogen (1:500) for 5h or 2′3′-cGAMP (25 μg/ml) for 18h along with Brefeldin A and Monensin. For CCL4 induction, Mitogen was used; conversely, 2′3′-cGAMP was used for CD40 and CD86 induction.

We titrated antibodies using freshly isolated and/or stimulated 270 μl of WB or 3×10^6^ of PBMCs per test (per MDIPA tube). All Abs were titrated in a two-fold dilution series started with a 1:50 dilution and titrated up to 1:200 dilution (Figure S2) according to the manufacturer’s (Standard BioTools) instruction ^[93]^.

#### Staining

CyTOF staining was performed according to the manufacturer’s (Standard BioTools) instruction ^[94]^. Briefly after stimulation, Fc blocked (Table S1) 270 μl of stimulated WB ^[95]^ or 3×10^6^ of PBMCs ^[96]^ were barcoded using metal-tagged CD45 Abs (110/111Cd and 194/195Pt) by using four-choose-three format ^[39]^ and added to MDIPA tubes for surface staining (30 minutes at RT). This approach of barcoding aims to minimize batch effects and reduce the sample acquisition time by CyTOF XT. After completion of surface staining, WB in each MDIPA tube was lysed using Cal-Lyse^TM^ Lysing solution (Table S1), according to the manufacturer’s (Standard BioTools) instruction ^[94–95]^. After RBC lysis (for WB only), barcoded clear cell pellets from four different conditions were combined (Figure S1A), washed and subjected to another round of surface stain using drop in Abs CD40, CD80, CD86 and CD69 (Table 1). The titer of each antibody used was 1:100. After completion of staining with drop-in activation markers, barcoded tubes were subjected to fixation (using 1.6% formaldehyde solution) (Table S1) followed by intracellular staining (30 minutes at RT) using a cocktail (1:100 dilution of each antibody in Maxpar^®^ Perm-S Buffer) (Table S1) of the functional cytokines (IFNγ, TNF, IL-2, IL-4, IL-17A), secretory proteins (Perforin, Granzyme B) and chemokine CCL4 (Table 1). After cell-ID^TM^ Intercalator (191/193Ir) staining (to identify nucleated cells), the stained cells were frozen at -80°C until acquisition by the CyTOF XT instrument.

#### Acquisition

During the day of acquisition, cells were washed with Maxpar^®^ Cell Staining Buffer (CSB) followed by Maxpar^®^ Cell Acquisition Solution (CAS) Plus, filtered through a 35μm cell strainer and pelleted. CyTOF XT calibration and data acquisition were performed according to the manufacturer’s (Standard BioTools) instruction ^[94]^. Briefly, a total of pelleted 1×10^6^ cells per tube were subsequently loaded onto the chilled carousel of the CyTOF XT at Cytometry Cores, situated within the Dana-Farber Cancer Institute. The CyTOF XT’s autosampler dispensed CAS Plus to the pelleted sample tubes, which were subsequently mixed with EQ^™^ Six Element Calibration Beads (Table S1) at a concentration of 0.1X. The CyTOF XT was then utilized to acquire the samples at a rate of 300-500 cells per second, employing CyTOF software (version 9.0.2). This acquisition process was accompanied by noise reduction, event length limitations set at 10-150 pushes, and a flow rate of 30μl/min.

### Analysis of CyTOF data

#### Quality control (QC)

After acquisition by the CyTOF XT instrument, data were normalized with CyTOF software (version 9.0.2). Normalized FCS files were de-barcoded manually (Figure S1B & C) using FlowJo (version 10.10) and uploaded to the cloud-based platform OMIQ (Dotmatics) ^[40]^ for further analysis. Upon reviewing the file metadata and file names by the OMIQ platform, scaling adjustments were made whenever the default settings were found unsuitable. After scaling adjustments, anomalous events were removed by the PeacoQC algorithm ^[97]^. For the PeacoQC run, the recommended settings in the OMIQ platform were followed (MAD: 6; Isolation Tree Gain Limit: 0.6; Minimum Number of Consecutive Bins: 5; Time Units Per Visualization Bin: 100 along with two selected parameters which were “Remove Zeros Before Peak Detection” along with “Generate Plots”). After the removal of anomalous events, quality-controlled FCS files were further cleaned up (Figure S1D) using recommended Gaussian parameters ^[98]^ using the OMIQ platform.

#### Manual gating

Following the data clean-up using Gaussian parameters, live nucleated cells (live 193Ir^+^) were manually gated further in accordance with MDIPA guidelines ^[99]^, which aligns with previous publications conducted by our group ^[29]^ and other researchers in the immunophenotyping realm ^[2, 32, 100]^. We adopted the gating strategy for phenotyping monocytes from OMIP-101 ^[101]^ and monocyte stages from OMIP-83 ^[102]^ (Figures 1A & S3A) since we did not follow the strategy of using CD38 for monocyte classification ^[32]^. We followed the MDIPA guideline for NK cell phenotyping ^[99]^. Gating CD123^-^ CD45RA^+^ cells during phenotyping of NK cell populations is not common. This specific gate ensures the purity of the NK cell population, as CD123 co-expression is observed in human NK cells derived from PBMCs ^[103]^ or vice versa ^[104]^. These observations justify the rationale for phenotyping NK cells as depicted in Figure 1A. Additionally, the stages of NK cells (Figure S3B), B cells (Figure S3C), Tregs (Figure S3D), and the memory phenotypes within the CD4^+^ (Figure S3E) and CD8^+^ (Figure S3F) T cell compartments were also mapped.

#### Quality assurance (QA)

Quality assurance is conducted by an independent consultant (Ionic Cytometry Solutions) to verify the file metadata, protocol, de-barcoding strategy, signal intensity of each marker in the CyTOF panel during data acquisition and the manual gating strategy by the OMIQ platform. Reference cryopreserved PBMCs (from a single donor) were used during different CyTOF runs and % CV was calculated to track down the deviation of cellular abundances (Figure S1E & Table S2). The assays exhibited a complete absence of technical variability because the % CV was below 25 (Table S2), which is a recommended criterion in deep immunophenotyping ^[105]^.

#### HD reduction analysis

A comparative study between tSNE-CUDA ^[42]^, opt-SNE ^[40]^ and FIt-SNE ^[41]^ were performed using the OMIQ platform to choose the most efficient HD reduction algorithm for downstream analysis. The Fit-SNE algorithm was selected for downstream analysis due to its ability to preserve the coordinates of cellular islands across different runs and its relatively faster execution compared to opt-SNE (Figure S4). For HD reduction analysis, 94K events of live CD45 population (CD45^+^193Ir^+^) were sampled equally per file using the “Subsampling” task by the OMIQ platform. FCS files are automatically excluded from the HD reduction and statistical analysis which does not possess 94K events of the chosen live CD45 population. We included all the 49 channels (Table 1) except LCBs (110/111 Cd and 194/195 Pt), 191/193 Ir and Live/Dead (103Rh) for DR analysis using tSNE-CUDA, opt-SNE and FIt-SNE (Figure S4).

For each tSNE-CUDA algorithm run, the parameters were Iterations = 1500, Early Exaggeration Iters = 250, Early Exaggeration Factor = 12, Perplexity = 50, Theta = 0.5, Learning Rate = 166,500, Num Nearest Neighbors = 32 and Print Progress Iters = 50.

For each opt-SNE algorithm run, the parameters were Max Iterations = 1500, opt-SNE End = 5000, Perplexity = 50, Theta = 0.5, Components = 2, Random Seed =7000 and Verbosity = 25.

For each FIt-SNE algorithm run, the parameters were Number of dimensions = 2, Perplexity = 50, Theta = 0.5, Random Seed =7000, Max iterations = 1500, Stop Early Exaggeration = 250, Learning Rate = 5000, Early Exaggeration Factor = 12.

After HD reduction analysis, all the desired cell populations from the cellular landscapes were overlaid in a scatterplot using the OMIQ platform. All the desired cell population abundances were calculated after “Subsampling” and graphed using GraphPad Prism version 10.4.2 for macOS for immunophenotyping.

#### CITRUS

ML algorithm CITRUS ^[48]^ was used by the OMIQ platform to dissect the stratifying signatures in the pDC compartment of PRRa stimulated PBMCs using the following default parameters: File Grouping: Stim; Min cluster Size Parcent: 0.05; Cross Validation Folds: 1; Regression Methods: pamr; Seed: 8610; Featuring Type: abundances. The CITRUS algorithm provided two predictive models (cv.min and cv.1se), each with varying stringency levels (Figure S6B). While a model with the absolute minimum error rate (cv.min) might be appealing, it can sometimes overfit the training data, meaning it performs very well on the data it was trained on but not so well on new data. The 1SE approach balances the desire for low error rate with the need for model simplicity, leading to a more robust and generalizable model. Therefore, the model cv.1se was selected for statistical output as it is the most predictive model for the exploratory dataset ^[106]^. The accuracy of the prediction model is presented in Figure S6B.

Before running CITRUS, DC populations were sampled equally by targeting 351 events of DCs per FCS files using the “Subsampling” task. The abundance of DCs was also manually calculated using the OMIQ platform (Figure 4B). The same strategy was employed to map the DC abundances in WB (Figure 4A) by employing equal sampling (284 events per FCS files). However, we refrained from utilizing CITRUS with DCs from WB due to the absence of PRRa-mediated mDC/pDC abundances when compared to steady-state conditions. After the CITRUS run (DCs from PBMCs), manual validation (to calculate p value) of cluster abundance was performed (Figure 4D) using categorical filters based on the cluster results in the OMIQ platform.

#### Statistics and reproducibility

Statistical significance and graphic output were generated using the GraphPad Prism (Dotmatics) version 10.4.2 for macOS. Data are represented in truncated violin plots as the median with interquartile range. Data were tested for normality using the Shapiro-Wilk test. The statistical analysis is meticulously documented in the figure legends. The sample size for *in vitro* stimulations was empirically determined based on our previous study ^[29]^. Furthermore, to mitigate batch effects, each CyTOF experiment utilized reference cryopreserved PBMCs and LCBs.

## Acknowledgements

D.J.D’s laboratory is funded in part by the U.S. National Institute of Health (NIH)/National Institutes of Allergy and Infectious Diseases (NIAID) Vaccine Adjuvant Discovery Program Contracts #75N93019C00044 and #75N93024C00020. The *Precision Vaccines Program* (PVP) is supported, in part, by Boston Children’s Hospital, Department of Pediatrics. The authors thank the members of the PVP and the Program Director Dr. Ofer Levy for their helpful discussions and support. The authors thank Suzan Lazo, Director of the Cytometry Cores at Dana-Farber Cancer Institute (DFCI) for her support and feedback. The authors also thank Eric Haas from Ionic Cytometry Solutions for assistance with data QA. S.B. would like to thank Emily M. Thrash (current employer: Lunaphore), Andrew Draghi II (current employer: Sony Biotechnology Inc.), Vinicius Motta (current employer: Nomic Bio) and Megan Perkins from Standard BioTools for their technical assistance and feedback on MDIPA Kit, MDIPA backbone and CD45 LCBs during the panel optimization. The authors extend their sincere gratitude to Dr. Justin Tirados of Dotmatics Ltd. for his invaluable assistance in the analysis of the data utilizing the OMIQ platform.

## Author Contributions

S.B. and D.J.D. conceived the project. Experiments were conducted by A.K. (Phlebotomy and PBMC isolation) and S.B. (stimulation and CyTOF stain). CyTOF acquisition by CyTOF XT was conducted by A.P. CyTOF data analysis, algorithm implementation, and presentation in figures were done by S.B. The manuscript was written by S.B. with feedback from A.P., A.K., and D.J.D. D.J.D. secured funding and supervised the study.

## Data Availability Statement

Data are available in the main article, figure, tables and in the supporting information. FCS files for CyTOF or other additional data which support the optimization of the study (i.e. titration and stained reference PBMCs) are available from the corresponding author or our *PVP* CyTOF team at upon reasonable request.

