## supporting information for "Deep Phenomics of Pattern Recognition Receptor Agonist Specific Activation of Human Blood"

### Table of Contents

| Supplementary Figures | | Page |
|---|---|---|
| Figure S1 | A unique LCB strategy, de-barcoding strategy, data clean-up using Gaussian parameters and criteria to avoid the high technical variability across the assay. | S-3 |
| Figure S2 | Optimal concentration of Abs is defined by titrations using freshly isolated WB and PBMCs. | S-5 |
| Figure S3 | Representative gating strategy to further classification of monocytes, NK cells, B cells, Treg, memory compartment of $CD4^{+}$ and $CD8^{+}$ T cells. | S-7 |
| Figure S4 | Evaluation of three different HD reduction algorithms tSNE-CUDA, opt-SNE and FIt-SNE using the OMIQ platform. | S-8 |
| Figure S5 | Tracking the abundance of canonical cellular lineages in WB and in PBMCs in steady-state. | S-10 |
| Figure S6 | Hierarchical clustering analysis conducted using CITRUS. | S-11 |
| Figure S7 | Tracking the CCL4 producing cellular lineages after PRRa stimulation. | S-13 |

| Supplementary Tables | | Page |
|---|---|---|
| Table S1 | Biological samples, reagents and algorithms for mass cytometry assay. | S-15 |
| Table S2 | The precision acceptance criteria for the CyTOF assay. | S-16 |
| Table S3 | The mean abundance of the targeted cellular populations in WB and PBMC compartments. | S-16 |

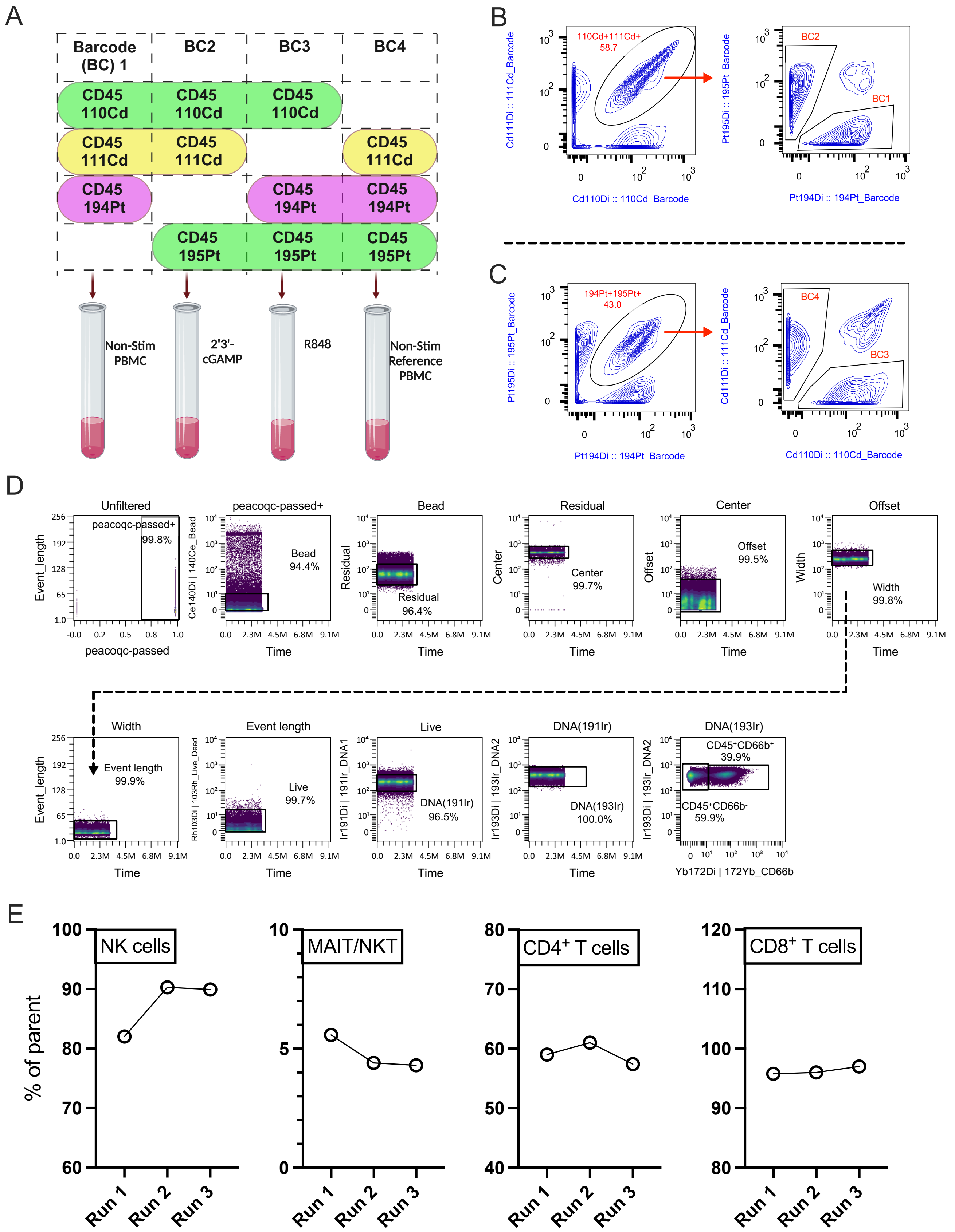

**Figure S1:** A) A unique LCB strategy employing the 4-choose-3 format was implemented, using an anti-CD45 barcode mixture (BC) which is compatible with MDIPA.

**Figure S1:** (Continued)

A representative example of four barcode sets (BC1 to BC4), each corresponding to a distinct stimulation, utilizing PBMCs. B, C) Gating strategy to de-barcode the CD45 cadmium (Cd) and platinum (Pt) tag using FlowJo 10.10. D) After de-barcoding, FCS files were subjected to PeacoQC algorithm using the OMIQ platform to remove anomaly. Quality-controlled (PeacoQC-passed) and assured CyTOF data were further cleaned up using the Gaussian parameter plotted against time to get out aggregates, debris, bead, doublet and dead cells. E) Assessing the abundance of representative major immune cell populations between CyTOF runs using 2′3′-cGAMP (25 µg/ml) stimulated cryopreserved reference PBMCs to measure the % CV. Such precision criteria were adopted and documented in **Table S2** to avoid the high technical variability across the assay.

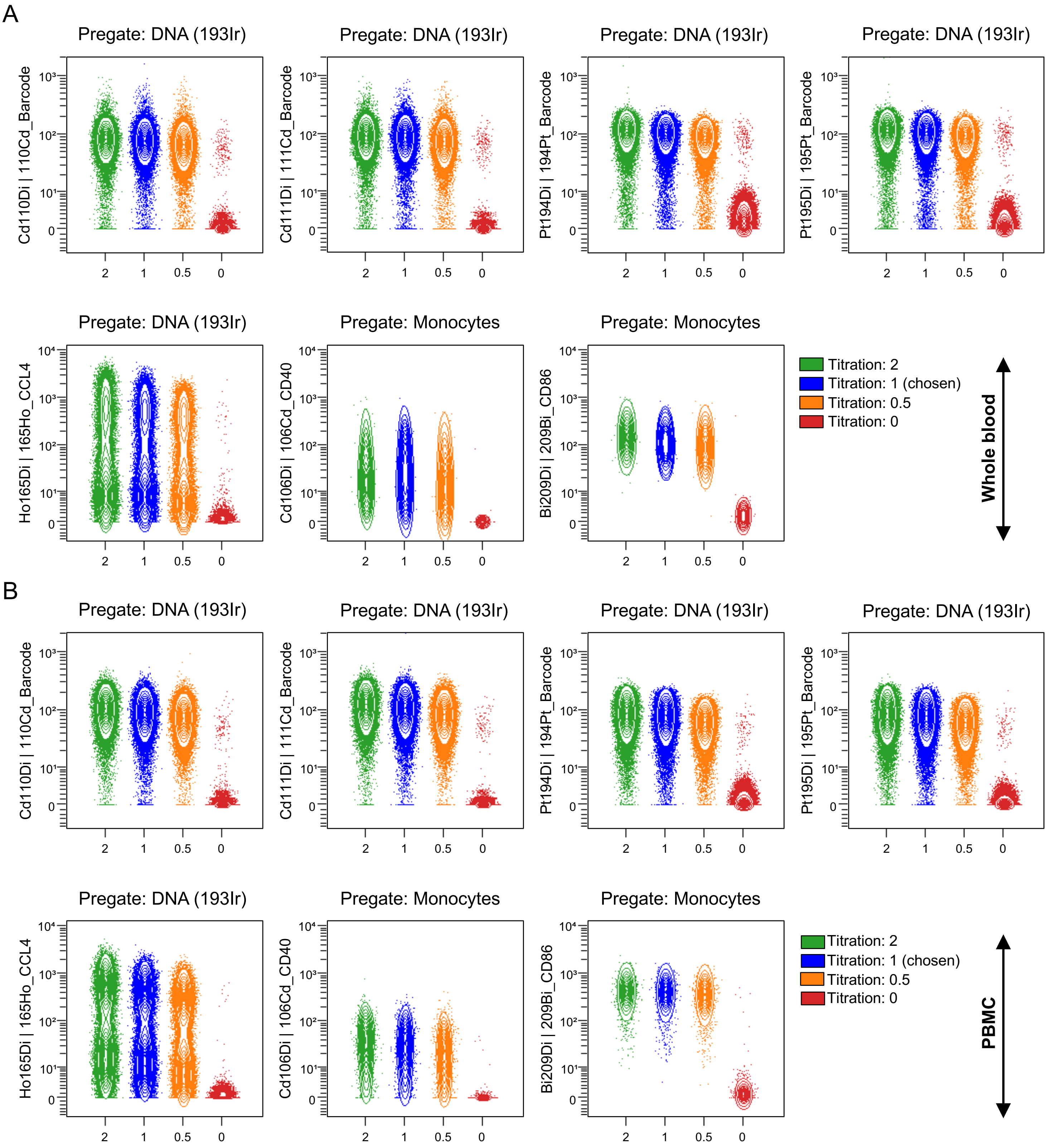

**Figure S2:** Optimal concentration of CD45 (110/111 Cd, 194/195 Pt) LCBs and custom-conjugated Abs (CCL4, CD40 and CD86) is defined by titrations using freshly isolated A) WB and B) PBMCs.

**Figure S2:** (Continued)

Serial fold-dilutions (on the X-axis) of the metal tagged Abs were started with 2 µL/100 µL, using mitogen (1:500) or 2′3′-cGAMP (25 µg/ml) stimulated cells to induce chemokine (CCL4) or activation markers and titrated thereafter. For LCBs and CCL4, WB and PBMCs were stimulated with mitogen, while 2′3′-cGAMP was used to induce activation markers CD40 and CD86 titrations. Cleaned up FCS files were gated to live $193Ir^{+}$ cells or monocytes and virtually overlaid according to the dilutions in the concatenated representation by the OMIQ platform for the representation. Blue subsets portray the chosen titer that provided the distinctive separation between positive and negative populations with minimal background staining.

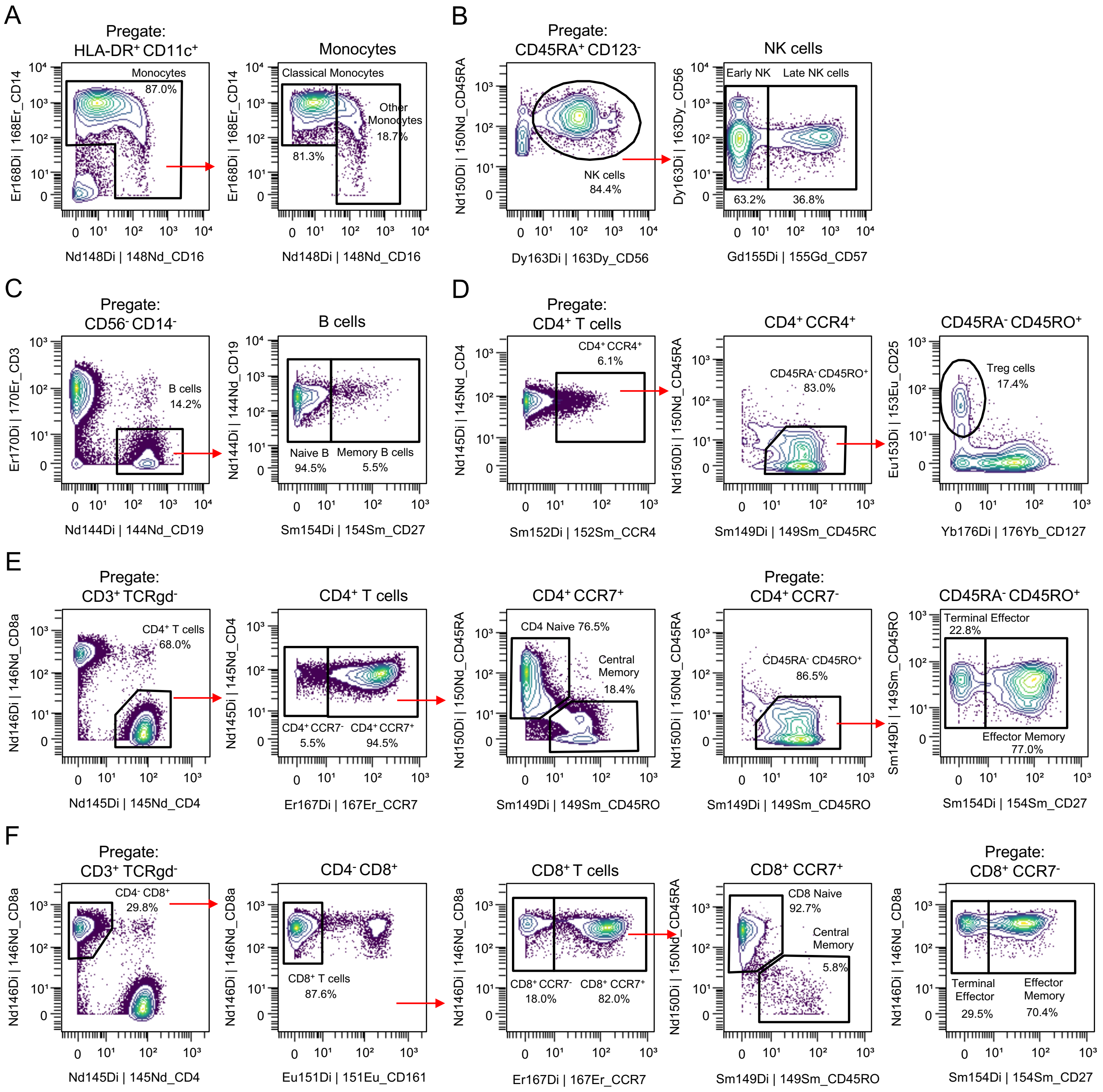

**Figure S3:** Representative gating strategy to further classification of (A) monocytes, (B) NK cells, (C) B cells, (D) $CD4^+$ T cells to define Regulatory T cells (Treg), memory compartment of (E) $CD4^+$ T cells and lastly (F) $CD8^+$ T cells. Red arrows represent sequential gating for selected populations. Manual gating was accomplished by the OMIQ platform.

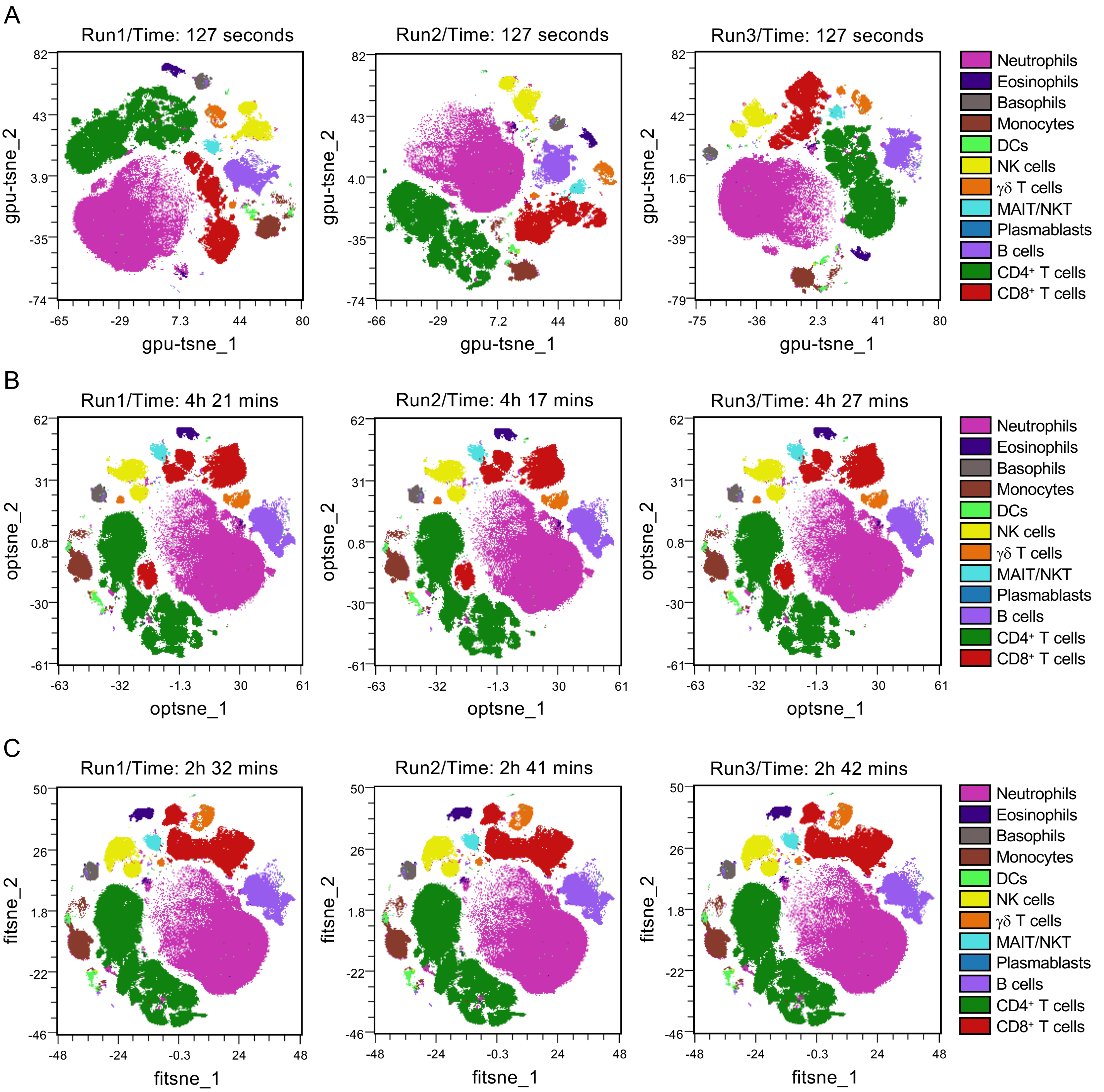

**Figure S4:** Evaluation of three different HD reduction algorithms (A) tSNE-CUDA, (B) opt-SNE and (C) FIt-SNE using the OMIQ platform in concatenated visualization. 94K events of live $CD45^+$ cells from each donor used in concatenated visualization with tSNE-CUDA or opt-SNE or FIt-SNE. Despite being the fastest (average run time of 127 seconds) HD reduction algorithm, tSNE-CUDA failed to maintain the coordinates of the cellular islands in the topography of tSNE-CUDA plots across multiple runs.

**Figure S4:** (Continued)

In contrast, opt-SNE and FIt-SNE demonstrated remarkable success in preserving the cellular island's coordinates and phenotypic distribution, thereby ensuring reproducibility of the cellular topography across different runs. For this study, the FIt-SNE algorithm was selected because it is the fastest, with an average run time of 2h 38 mins, compared to the opt-SNE algorithm, which has an average run time of 4h 21 mins using the same data set and parameters.

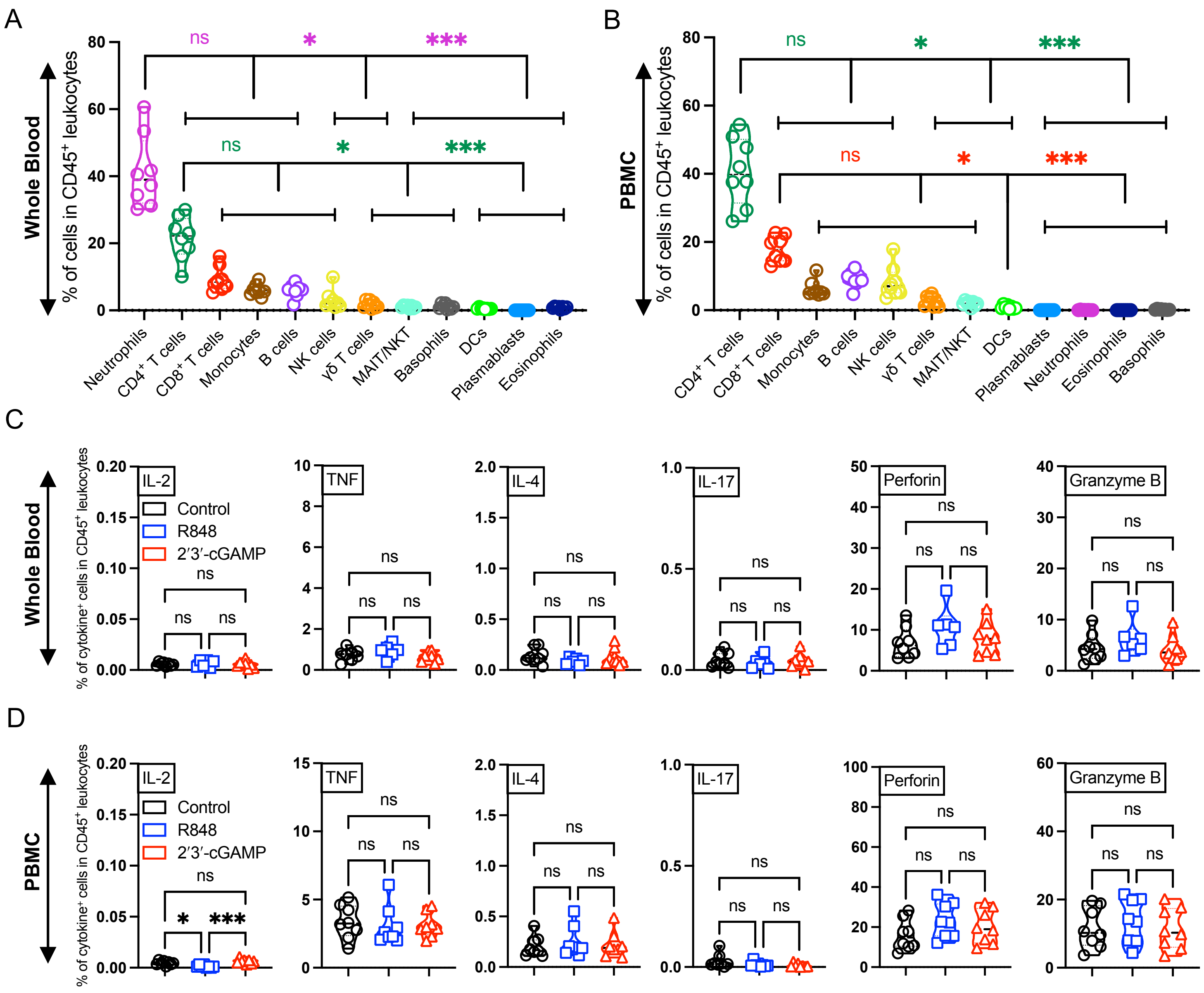

**Figure S5:** Tracking the abundance of canonical cellular lineages in (A) WB and in (B) PBMCs in steady-state. The threshold criteria for the downstream analysis were set as 94K events of live CD45⁺ cells from each donor, as described in Figure 1. C, D) Intracellular cytokines (IL-2, TNF, IL-4, IL-17) and secretory proteins (Perforin, Granzyme B) profile of live CD45⁺ cells after PRRa stimulation for 18h. C) WB or (D) PBMCs stimulated with R848 (5μM) and 2′3′-cGAMP (25 μg/ml) using the adult cohort. Statistical comparison was performed using either one-way ANOVA or nonparametric Kruskal-Wallis test corrected for multiple comparisons; *p < 0.05, ***p < 0.001, ns denoted non-significant. Each circle, square or triangle represents a single participant (n=6-8 per group).

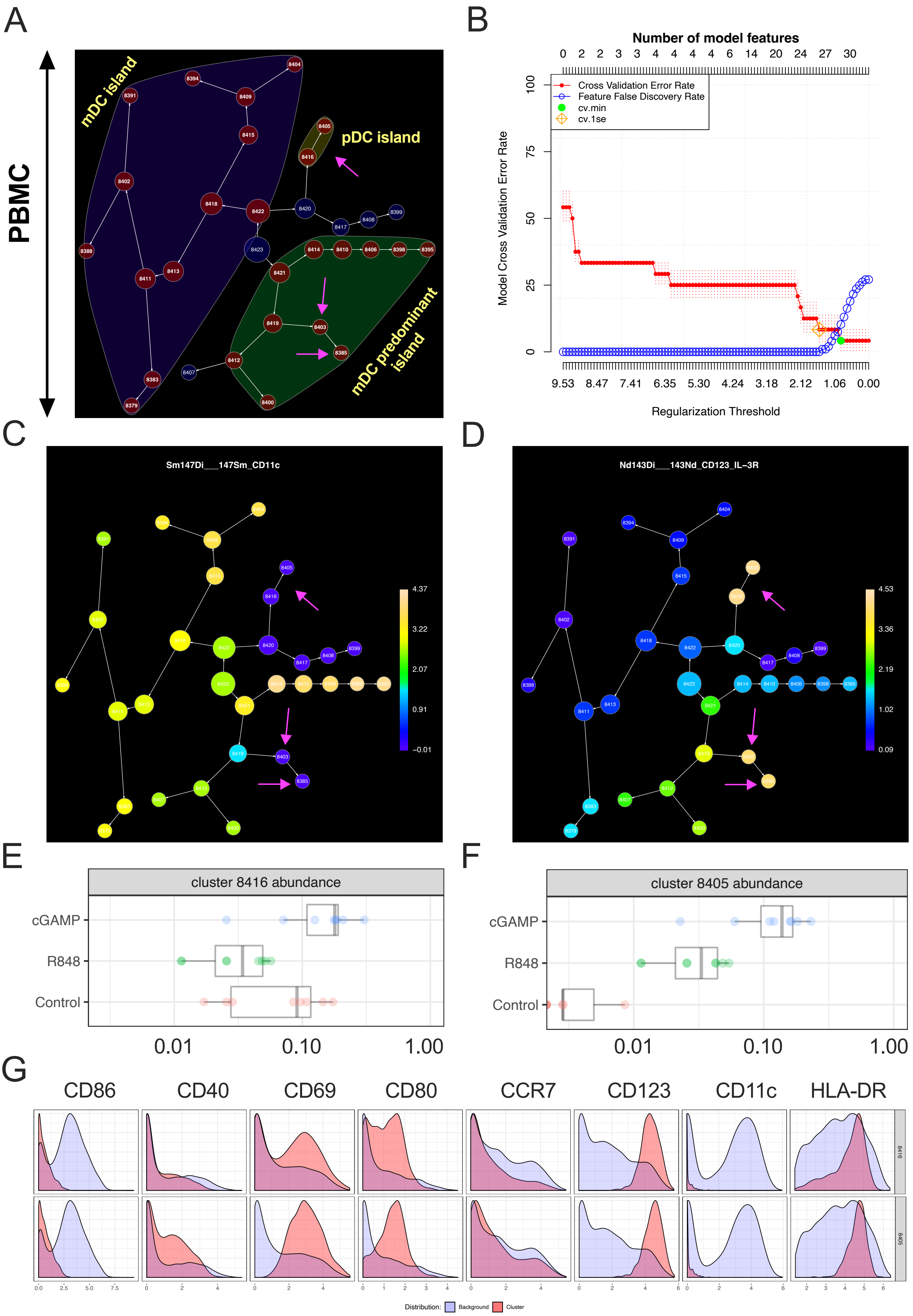

**Figure S6:** A) Hierarchical clustering analysis conducted using CITRUS was employed to stratify the subpopulations of DC based on HLA-DR, CD11c, CD123, CCR7, CD80, CD69, CD40, and CD86 markers (Table 1).

**Figure S6:** (Continued)

Maroon clusters showed significant differences, while blue clusters lack them. The numerical value within each circle represents the cluster ID. White arrows between clusters illustrate the relationship, while magenta arrows indicate the pDC clusters of interest. B) Accuracy of the prediction model. The cv.1se predictive model (orange square with two diagonals) was chosen for cluster abundance and phenotype prediction. The cv.1se model showed low cross validation error rate (red line) along with low FDR (blue line). C) CD11c and (D) CD123 expression on DCs after hierarchical clustering of PBMCs. Color scales illustrate the relative (C) CD11c and (D) CD123 expression per cluster while size of each node represents event frequency. Based on the relative expression of CD11c and CD123, we named the cluster islands in (A). Graphs generated by CITRUS showed the quantitative results of abundance from E) cluster 8416 and F) cluster 8405 between PRRa stimulated groups and steady-state. Phenotypic expression of these clusters is also shown in G). The expression level of each selected marker was represented by a cluster (red) over its background (light blue). Figures (A-G) were directly exported from the OMIQ platform.

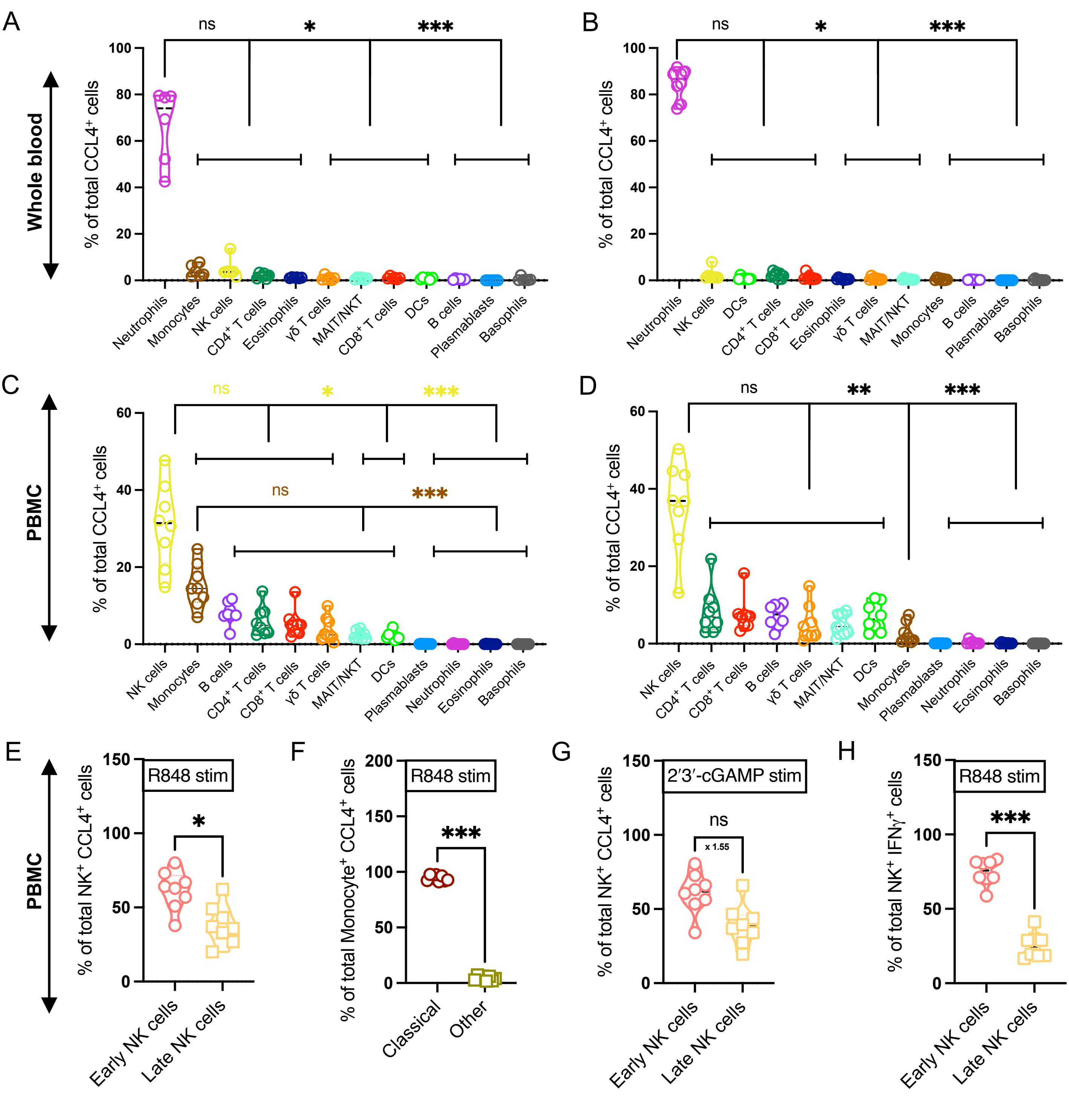

**Figure S7:** Tracking the CCL4 producing cellular lineages after PRRa stimulation. (A) CCL4 production by major immune cell subsets in WB after R848 (5µM) or (B) 2′3′-cGAMP (25 µg/ml) stimulation for 18h. CCL4 production was also captured in (C) PBMC after R848 or (D) 2′3′-cGAMP stimulation.

**Figure S7:** (Continued)

(E) CCL4 producing NK cell and (F) monocyte compartment was dissected to gain more insight about CCL4 producing cellular lineages after R848 stimulation. Further profiling of (G) 2′3′-cGAMP mediated CCL4 production and (H) R848 mediated IFNγ production in NK cell compartment. Mean fold difference between early and late NK cells is also shown after 2′3′-cGAMP stimulation. Statistical comparison was performed using either one-way ANOVA or nonparametric Kruskal-Wallis test corrected for multiple comparisons. For E-H, two-tailed paired t-test was used for comparison between 2 groups; *$p < 0.05$, **$p < 0.01$, ***$p < 0.001$, ns denoted non-significant. Each circle or square represents a single participant (n=6-8 per group).

**Table S1.** Biological samples, reagents and algorithms for mass cytometry assay.

| Reagent or Resource | Source | Identifier |
|---|---|---|
| Biological samples | | |
| Blood from healthy adult (22-33) participants were collected between August and September 2023 | --- | --- |
| Chemicals, TLR agonists | | |
| Ficoll-Paque™ PREMIUM | Cytiva | 17544203 |
| Resiquimod (R848) (TLR7/8a) | InvivoGen | tlrl-r848-1 |
| 2′3′-cGAMP (STINGa) | InvivoGen | tlrl-nacga23-1 |
| Maxpar® Direct™ Immune Profiling Assay (MDIPA) Kit | Standard BioTools | 201334 |
| Maxpar® Direct™ T Cell Activation Expansion Panel | Standard BioTools | 201409 |
| Cal-Lyse™ Lysing Solution | Invitrogen | GAS010 |
| Maxpar® Water | Standard BioTools | 201069 |
| Pierce™ 16% Formaldehyde | Thermo Scientific | 28906 |
| Maxpar® Perm-S Buffer | Standard BioTools | 201066 |
| Heparin Sodium Injection, 10KU/10ml | Meitheal Pharmaceutical | 71288-402-11 |
| EQ™ Six Element Calibration Beads | Standard BioTools | 201245 |
| Brefeldin A Solution | BioLegend | 420601 |
| GolgiStop (containing Monensin) | BD Biosciences | 554724 |
| Human BD Fc Block™ | BD Biosciences | 564220 |
| Cell Activation Cocktail | BioLegend | 423301 |
| Software and Algorithms | | |
| FlowJo (10.10) for macOS | BD | https://www.flowjo.com/flowjo/download |
| OMIQ (accessed between January and May 2025 for this study) | Dotmatics | https://www.omiq.ai |
| PeacoQC | Dotmatics | https://www.omiq.ai |
| tSNE-CUDA, opt-SNE, FIt-SNE | Dotmatics | https://www.omiq.ai |
| CITRUS | Dotmatics | https://www.omiq.ai |
| Prism 10 (Version 10.4.2) for macOS | GraphPad Software | https://www.graphpad.com |
| EndNote 21 | Clarivate | https://endnote.com |
| Microsoft 365 | Microsoft | https://www.microsoft.com/en-us/microsoft-365 |
| BioRender | BioRender | https://www.biorender.com |

| | NK cells | Early NK | Late NK | DCs | Monocytes | Classical Mono | Other Mono | MAIT/NKT |
|---|---|---|---|---|---|---|---|---|
| Run 1 | 82 | 65.7 | 34.3 | 7.39 | 38.1 | 97.1 | 2.73 | 5.58 |
| Run 2 | 90.3 | 64.5 | 35.5 | 6.62 | 30.4 | 95.5 | 4.26 | 4.4 |
| Run 3 | 89.9 | 66.6 | 33.4 | 9.67 | 30.4 | 96.9 | 3.07 | 4.3 |
| Mean | 87.40 | 65.60 | 34.40 | 7.89 | 32.97 | 96.50 | 3.35 | 4.76 |
| SD | 3.82 | 0.86 | 0.86 | 1.30 | 3.63 | 0.71 | 0.66 | 0.58 |
| % CV | 4.37 | 1.31 | 2.50 | 16.41 | 11.01 | 0.74 | 19.56 | 12.21 |

| | $CD4^+$ T cells | $CD8^+$ T cells | $T_{reg}$ | TCRγδ | B cells | Plasmablasts | $DNA^+$ $CCL4^+$ | $DNA^+$ $IFN\gamma^+$ |
|---|---|---|---|---|---|---|---|---|
| Run 1 | 59 | 95.8 | 15.9 | 31.3 | 21.6 | 17.4 | 9.24 | 0.306 |
| Run 2 | 61 | 96 | 19.3 | 43.4 | 29.5 | 12.5 | 11.5 | 0.313 |
| Run 3 | 57.4 | 97 | 13.6 | 33.1 | 21.6 | 10.3 | 8.56 | 0.234 |
| Mean | 59.13 | 96.27 | 16.27 | 35.93 | 24.23 | 13.40 | 9.77 | 0.28 |
| SD | 1.47 | 0.52 | 2.34 | 5.33 | 3.72 | 2.97 | 1.26 | 0.04 |
| % CV | 2.49 | 0.55 | 14.39 | 14.83 | 15.37 | 22.15 | 12.87 | 12.56 |

**Table S2:** The precision acceptance criteria for the CyTOF assay. The percent coefficient of variance (% CV) for each population (as % of parent) were measured to determine the technical variation across batches. For each assay, reference control PBMC were stimulated using 25µg/mL of 2′3′-cGAMP and analyzed by calculating the mean abundance, standard deviation (SD), and % CV. A high technical variable was absent from the assays, as the acceptance criterion was set at a CV of less than or equal to 25%.

| | Neutrophils | $CD4^+$ T cells | $CD8^+$ T cells | Monocytes | B cells | NK cells | Early NK | Late NK |
|---|---|---|---|---|---|---|---|---|
| WB | 41.19 (***) | 21.53 | 9.355 | 6.099 | 5.871 | 3.145 | 2.114 | 1.031 |
| PBMC | 0.0694 | 40.73 (***) | 17.85 (***) | 6.233 | 9.166 (***) | 8.284 (**) | 4.929 (*) | 3.358 (**) |

| | γδ T cells | MAIT/NKT | Basophils | DCs | Plasmablasts | Eosinophils | $T_{reg}$ |
|---|---|---|---|---|---|---|---|
| WB | 1.716 | 1.058 | 1.179 (**) | 0.5616 | 0.0092 | 0.8385 (***) | 0.1185 |
| PBMC | 2.610 (**) | 1.973 (**) | 0.1917 | 0.9074 | 0.0097 | 0.0205 | 0.4675 (***) |

**Table S3:** The mean abundance of the targeted cellular populations in WB and PBMC compartments (as described in Figure 1 and Figure S5) is also shown in a tabular format. The downstream analysis was conducted with a threshold of 94K live $CD45^+$ cells from each donor (n = 8 per group). Statistical comparison was performed using either two-tailed paired t-test or Wilcoxon test corrected for multiple comparisons; *p < 0.05, **p < 0.01, ***p < 0.001. Statistical significance between groups is also highlighted by a light green color.
